# Emotional amnesia in humans with focal temporal pole lesions

**DOI:** 10.1101/2025.10.22.683695

**Authors:** Robin Hellerstedt, Manuela Costa, Dominic Giles, Diana Ortega-Cruz, Alba Peris-Yagüe, Lukas Imbach, Johannes Sarnthein, Debora Ledergerber, Philip Grewe, Johanna Kissler, Friedrich G. Woermann, Christian G. Bien, Antonio Gil-Nagel, Rafael Toledano, Bryan A. Strange

**Author notes:** Address for correspondence: Robin Hellerstedt and Bryan Strange.

## Abstract

Memory is typically better for emotional relative to neutral events, a process involving amygdala modulation of hippocampal activity^1–6^. These structures, however, form part of a larger emotional brain network, which in humans includes the temporal pole^7^, a cortical node whose functional role in emotional cognition remains poorly understood^8–10^. Here, we show, in pharmaco-resistant epilepsy patients performing verbal and visual emotional episodic memory tasks, a selective impairment in recalling verbal emotional memories in left ventral temporal pole (vTP) lesioned patients compared with control patients. Memory for neutral words, and verbal comprehension performance on standard neuropsychological testing, were intact in these patients, indicating absence of general episodic memory or semantic impairment. All patients underwent recordings with intracranial electrodes during memory task performance. Left vTP lesioned patients showed no differences in amygdala or hippocampal electrophysiological responses to emotional words, compared with control patients, putatively isolating a vTP role in emotional memory. Unlike verbal emotional recall, left vTP lesioned patients showed memory enhancement for emotional vs. neutral pictures, whereas two patients with right vTP lesions showed the opposite pattern: impaired memory for emotional pictures but intact verbal emotional memory. These observations establish a critical, lateralized, modality-specific role for human vTP in emotional memory, imply emotional memory deficits in neurological conditions affecting this region, and advance the vTP as a target for neuromodulation in diseases characterized by maladaptive emotional memories.

## Introduction

Better memory for emotionally negative relative to neutral events is evolutionary adaptive as it helps us to avoid previously encountered threats or aversive situations. Human lesion studies have established that, while hippocampal lesions impair memory for both emotional and neutral episodes, amygdala lesions selectively abolish emotional episodic memory enhancement, rendering negative emotional memory performance equivalent to that of neutral memory^2,3^. Functional magnetic resonance imaging (fMRI) studies^4,5,11^, and more recently human intracranial electrophysiological studies^6^, support the memory modulation hypothesis^1^ that amygdala activity modulates hippocampal function during emotional memory formation. By contrast, emotional memory retrieval is associated with a shift to a more hippocampal-centred mechanism^4,12^.

However, the functional coupling between amygdala and hippocampus forms part of a larger emotion-sensitive brain network^7^. The temporal pole (TP) was contemplated as a key cortical component of this network following the description of loss of emotion reactivity in Klüver-Bucy syndrome in non-human primates^13^ and later humans^14^, yet its contribution to emotional cognition remains poorly understood. Anatomically, the TP shows dense connectivity with the amygdala and the orbitofrontal cortex and has traditionally been included within the paralimbic system^15^. Although the anterior temporal lobe, including the TP, is considered a key amodal hub in the semantic network^16^, recent models have highlighted the importance of the TP in emotional and social-semantic processing relative to other subregions of the anterior temporal lobe^17^. The TP is implicated in social processes such as theory of mind and person memory (i.e. semantic memory for people’s identity, names, biography, traits etc.), via a functional role in encoding and retrieval of socio-emotional concepts^9,10,18^. In line with TP involvement in emotional processes, patients with the temporal variant of frontotemporal dementia and predominantly right sided atrophy show emotion-related symptomatology including depression and reduced empathy^19,20^. Furthermore, TP volume reductions have been observed in patients with psychiatric disorders that affect emotional processing including major depressive^21^ and bipolar^22^ disorders. Early PET studies in patients with post-traumatic stress disorder (PTSD) showed increased temporal polar responses during script-driven imagery symptom provocation^23,24^, and recent lesion network mapping implicates the anterior temporal lobe in a network where damage reduces PTSD risk^25^.

In the context of emotional memory, the TP has been proposed to reflect the psychological set associated with emotional memory retrieval^26^. However, a difficulty in studying the role of this structure in emotional memory (and in cognition in general) is that conditions producing TP lesions typically extend into other parts of the temporal lobe^27^, such as temporal variant frontotemporal dementia, which also produces hippocampal and amygdala damage^28^, or herpes-simplex encephalitis, which also affects most of the limbic areas^29^. In addition, susceptibility artefacts on fMRI due to air-tissue interfaces with the adjacent paranasal sinuses make imaging the TP challenging, unless sequences are optimized for studying this area, so it is likely that fMRI-studies have generally underestimated the role of TP in cognitive functions^30^.

In a series of medication-resistant epilepsy patients performing emotional memory tasks whilst undergoing intracranial recordings, we identified five patients with focal left ventral TP (vTP) damage (of whom one had bilateral vTP damage), and one patient with unilateral focal right vTP damage. We report here specific emotional memory deficits observed in all studied patients with vTP lesions. In a test of verbal episodic memory, left vTP lesioned patients showed a selective deficit for recalling infrequent emotionally negative words, but intact memory enhancement for infrequent, perceptually salient words. In contrast to previously described effects of amygdala lesions, which produce equal memory for emotional and neutral items^2,3^, left vTP lesioned patients recalled *fewer* emotional than neutral items. In a test of visual memory for emotionally negative and neutral pictures, however, memory for negative pictures in left vTP lesioned patients was at the same level as non-TP-lesioned patients. By contrast, the two patients with right vTP lesions showed worse memory for negative vs neutral pictures.

## Results

Five patients were identified with left vTP lesions: two with focal dysplasia, one due to previous trauma, and two with encephalocele. One of these 5 patients had bilateral TP encephaloceles, and a sixth additional patient had a right unilateral TP encephalocele (**Fig. 1a-b,** the aetiology and lesion location of all patients are listed in **Supplementary Table 1**). Note that lesions were highly focal to the TP, except for a left vTP patient with traumatic aetiology with more extensive damage, including the left lateral and inferior temporal gyrus. Lesion mapping showed overlap in the anterior tip of Brodmann areas 20,36 and 38 on the ventral aspect of the temporal pole, with all lesions overlapping with areas labelled “*Temporal Pole”* on the Atlas of Intrinsic Connectivity of Homotopic Areas^31^ (AICHA). The percentage of overlap with Brodmann areas and AICHA areas for each lesion are presented in **Extended Data Fig. 1-2.** A further 26 patients with treatment resistant epilepsy, undergoing depth recordings but without lesions in the temporal pole, were assigned to the control group (**Supplementary Table 1**).

**Fig. 1.**
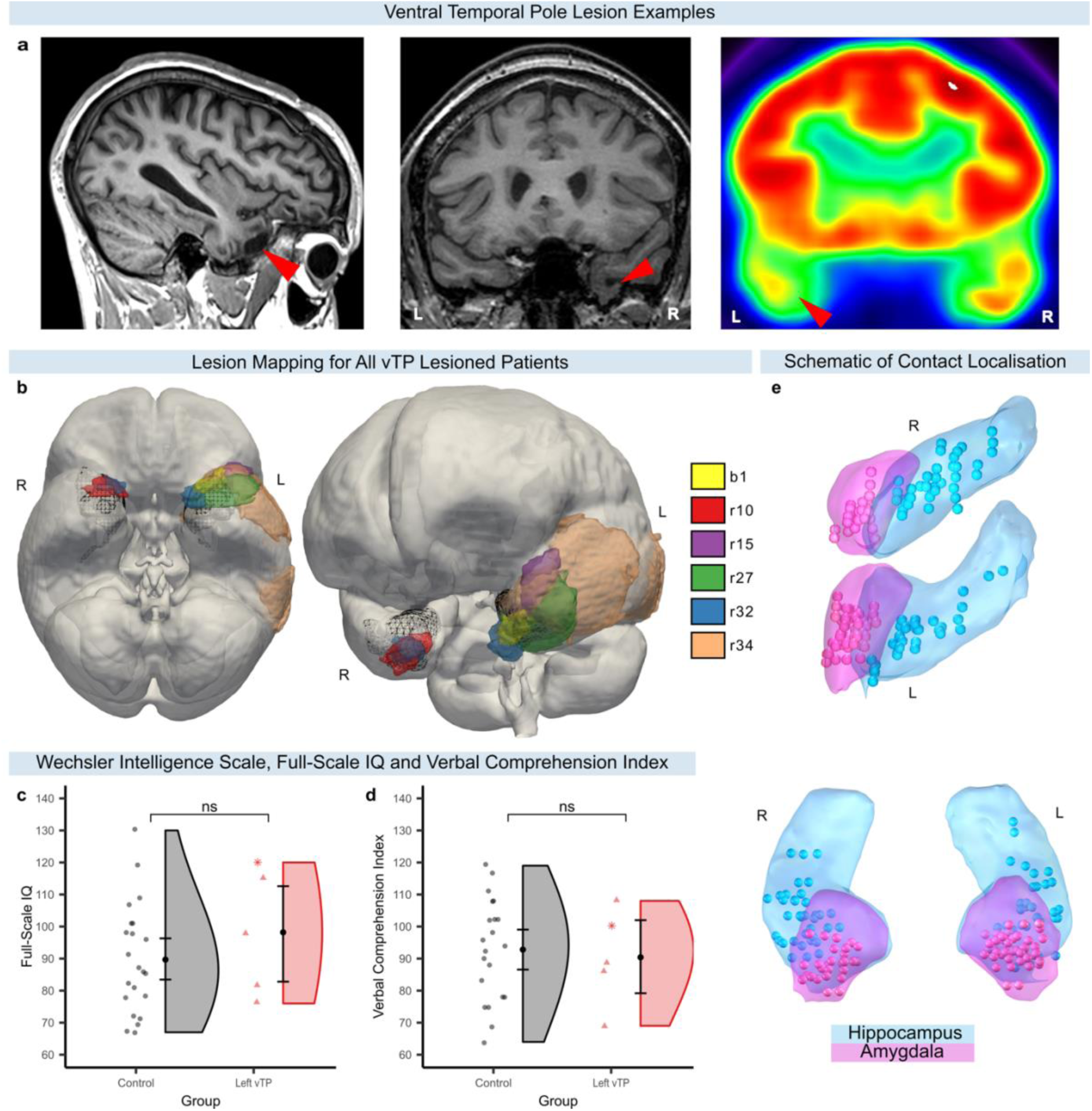
Example of temporal pole lesions, schematic of iEEG contact localization and neuropsychological test performance. **a**, Left, T1-weighted MRI example of a structural left vTP lesion due to traumatic brain injury (patient r34); Middle, T1-weighted MRI of right vTP lesion due to TP encephalocele (patient r10); Right, Fluorodeoxyglucose positron emission tomography (FDG-PET) example of a functional left vTP lesion due to focal dysplasia (patient r15; red colours indicate higher FDG uptake). **b**, Mapping of vTP lesions in MNI space. The TP area is depicted as a black mesh while colours indicate each patient’s lesion. **c**, Full-scale IQ from Wechsler intelligence scale tests for the control group and the left vTP group. Scatter dots display individual scores (circle = control, triangle = unilateral left vTP lesion, star = bilateral vTP lesions), black dots indicate group means and the error bars represent 95% confidence interval of the mean. **d**, Verbal comprehension index scores from Wechsler intelligence scale tests. **e**, Intracranial EEG contact localization in the left and the right amygdala (pink) and the hippocampus (light blue) for all patients included in both the verbal and the visual task. Amygdala: 22 patients, 27 electrodes, 74 contacts; Hippocampus: 22 patients, 28 electrodes, 68 contacts (all contacts are displayed on thresholded, post-operative CTs overlayed on a pre-operative MRI in **Supplementary Fig. 1**).

On neuropsychological testing prior to electrode implantation, the left vTP patients (*n* = 5) showed no difference to the control group (*n* = 23 for IQ, *n* = 21 for verbal comprehension) in full-scale IQ (*U* = 40.5, *P* = .322) or the verbal comprehension index (*U* = 58.5, *P* = .720) (**Fig. 1c-d**). The unilateral right vTP patient had an IQ and verbal comprehension score within the normal range (full-scale IQ = 97, verbal comprehension score = 100). Electrode contact localization in amygdala and hippocampus across all patients is shown in **Fig. 1e**. Electrode placement was based strictly on clinical grounds, thus not all patients had electrodes in both amygdala and hippocampus. Furthermore, not all patients participated in both verbal and visual tasks (**Supplementary Tables 1-7** list which patients were included in each analysis).

### Focal left ventral temporal pole lesions impair emotional verbal memory

In the verbal free recall task, the 22 participants (4 left vTP patients, 1 bilateral vTP patient, 1 right vTP patient and 16 control patients) saw 14 semantically-related nouns presented on-screen one at a time. In each list, one word was emotionally negative, and one was presented in a different font (perceptual oddball; **Fig. 2a**). The participant’s memory for the encoded words was tested by self-paced free recall following a brief distractor task after each list (**Fig. 2b**). All patients performed the Spanish version of this task, except for one left vTP patient, who performed the German version.

**Fig. 2.**
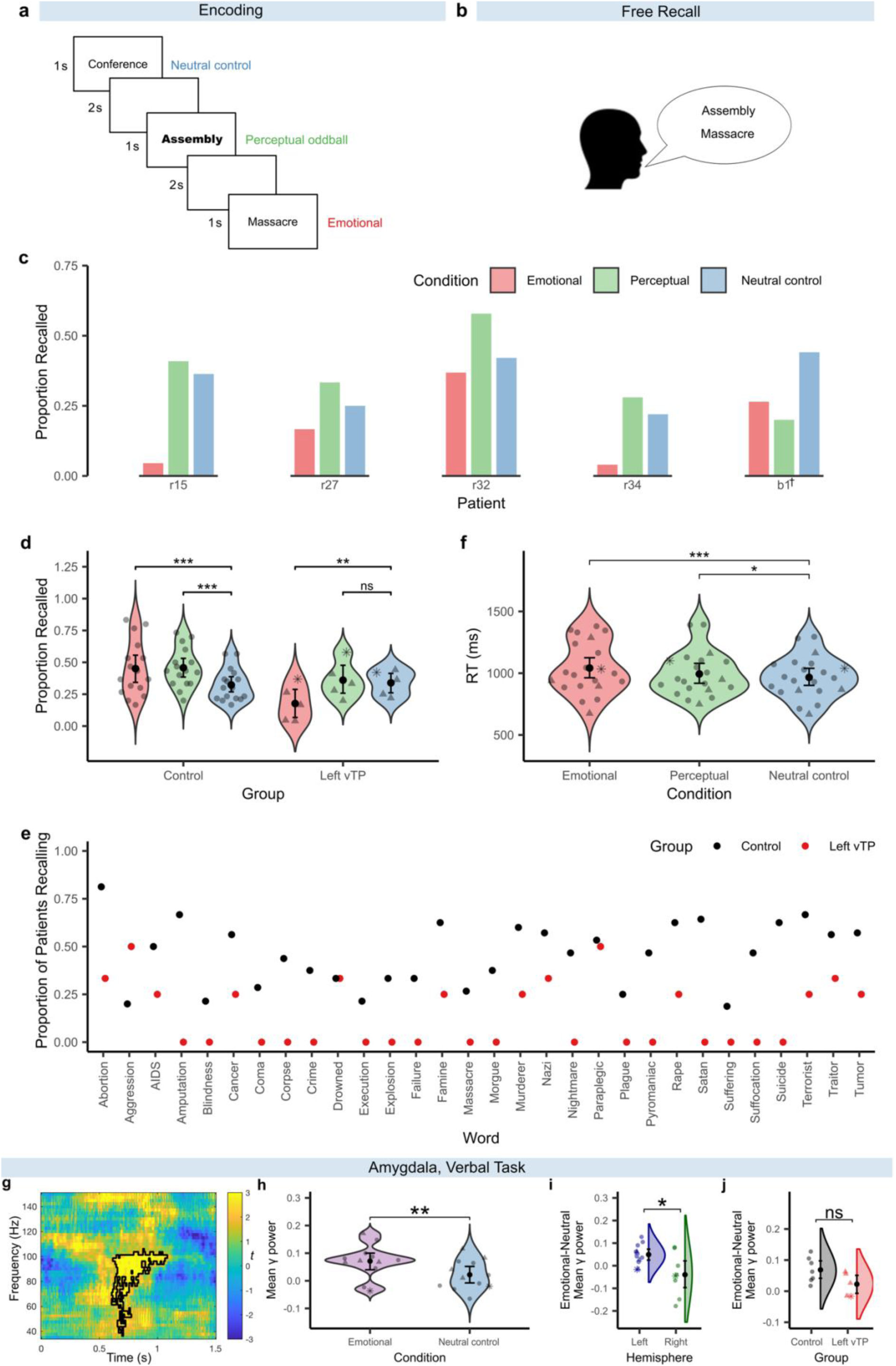
Experimental design, behavioural results and amygdala intracranial recordings in the verbal emotional memory task. **a**, Encoding phase of the verbal emotional memory task with examples of the three word-types: neutral control words, emotionally negative word and perceptual oddball (written in a different font). **b**, A self-paced free recall test was given after the encoding of each list. **c**, Memory performance for each of the five patients with left vTP lesions separated by condition. Note, patient r32 has bilateral vTP lesions. †There was a computer failure affecting the change of font in the perceptual oddball condition for patient b1. **d**, Memory performance in the free recall test separated by Group and Condition. Here, and in subsequent panels, scatter dots represent individual means (• control patient, ▴ unilateral left temporal pole lesion, * bilateral temporal pole lesion), the black dot indicates the group mean and the error-bars indicate the 95% confidence interval of the mean. Stars indicate significance levels (*** < .001, ** < .01, * < .05, ns = not significant). **e**, Proportion of patients in the control group and in the left temporal pole group remembering each of the 30 emotional oddball words. **f**, Reaction times in the encoding task (in which the participants judged if the initial letter contained a closed space) for each condition. **g,** Time-resolved spectral power induced by word presentation (time 0ms) in left amygdala. Colour scale indicates *t*-values from the comparison of emotionally negative vs neutral words, with black outline indicating a significant cluster from cluster-based permutation tests. **h**, Extracted mean gamma power from the significant cluster. Scatter dots display individual means, black dots indicate group means and error bars represent 95% confidence interval of the mean. Significance levels: ** < .01, * < .05, ns = not significant. **i**, Mean gamma power from the significant cluster from the left amygdala is now also shown for the right amygdala. **j**, Mean gamma power from the significant left amygdala cluster separated by control and left vTP groups.

Whereas verbal memory for neutral control words was not different between vTP-lesioned patients and control patients, a striking pattern was evident for emotional nouns. All 5 patients with left vTP lesions showed lower memory performance for emotional compared with neutral words (**Fig. 2c**). This pattern was only observed in 3 out of 16 (18.8%) of the patients in the control group and was absent in the only patient with unilateral right vTP lesion. We analysed verbal memory performance using a binomial generalized linear mixed-effects model (GLMM) with fixed effects of Group (control vs left vTP) and Condition (emotional oddball vs perceptual oddball vs control words) and with patient ID as a random effect with random intercepts (**Fig. 2d**). This random effect structure was used in all subsequent GLMM analyses described below. We observed a significant Group × Condition interaction (χ^2^(2) = 23.62, *P* < .001) and a main effect of Condition (χ^2^(2) = 9.61, *P* = .008), without a significant main effect of Group (χ^2^(1) = 3.67, *P* = .055). Follow-up pairwise comparisons showed that, in the control group, memory was higher for emotional words (*z* = 4.89, *P* < .001, odds ratio (OR) = 1.81) and for perceptual oddballs compared with neutral control words (*z* = 4.81, *P* < .001, OR = 1.80). No difference in memory between emotional and perceptual items was observed (*z* = .07, *P* = .945, OR = 1.01). In contrast, the left vTP lesioned group showed reduced memory for emotional words compared with neutral control words (*z* = −3.12, *P* = .003, OR = .43) and compared with perceptual oddballs (z = −3.11, *P* = .003, OR = .38). A second set of follow-up pairwise comparisons showed that while the vTP lesioned patients had reduced memory for emotional words compared with the control patients (*z* = −3.72, *P* < .001, OR = .26) there were no group differences in memory performance for neutral control words (*z* = 0.32, *P* = .748, OR = 1.10) and perceptual oddballs (*z* = −1.13, *P* = .257, OR = .68).

### Emotional word amnesia is unlikely to be due to other word properties

Next, we tested whether other aspects of presented emotional words, known to affect episodic memory, could explain the observed amnesia for these stimuli. Memory is enhanced for frequent words^32^, for words with high semantic associative strength to the other words in the list^33^, for concrete words^34^ and for words that more easily evoke a mental image (imageability)^35^. Of relevance for the present study, patients with anterior temporal lobe lesions have been shown to have larger impairments in semantic tasks (e.g. synonym judgement) for words that are less frequent and less imageable^36^.

In a first analysis we added semantic association strength between each word and the general list context (based on ratings), and word frequency per million words in a corpus (EsPal^37^, a repository with a comprehensive set of Spanish word properties based on texts from several sources on the web including news articles, literature and academic texts) as covariates in the GLMM with fixed effects of Group (control *n =* 16 vs left vTP lesioned *n* = 5) and Condition (emotional words vs perceptual oddball vs control words) described above. Importantly, the Group × Condition interaction remained significant (χ^2^(2) = 19.67, *P* < .001) after including these covariates. A second analysis included subjective ratings of word familiarity, imageability and concreteness (obtained from the EsPal database^37^ ) as covariates. The reason for conducting a separate analysis for these covariates was that these latter measures were not available for all words included in our study. The Group × Condition interaction remained significant in this model as well (χ^2^(2) = 9.39, *P* = .009). This interaction was also significant when including the German-speaking patient in the model with all five covariates (χ^2^(2) = 25.70, *P* > .001), even though the German word measures were not fully equivalent to the Spanish measures (see Methods).

In addition, we performed an item analysis to investigate if the memory impairment for emotional words was driven by specific words. This descriptive analysis showed that 28 out of the 30 negatively valenced words were remembered by a lower proportion of the subjects in the left vTP lesioned group than in the control group, indicating that the impairment was general (**Fig. 2e**).

### Focal left temporal pole lesions do not affect reaction times to emotional words

A possible explanation for the relative amnesia for emotional words in left vTP lesioned patients is that they were agnostic to their aversive meaning during encoding. One way to probe this is via reaction times to emotional vs neutral words. During word list presentation, patients made push-button responses to each word to indicate whether the initial letter contained a closed space (a “shallow” encoding task, designed to minimize elaborate semantic processing^38^). Healthy human adults show slower reaction times on this task for emotional words and perceptual oddballs compared with control words^2,39^. Here, we observed that both control patients and left vTP lesioned patients showed the expected slowing in response times to oddballs (**Fig. 2f, see Supplementary Data 1 for statistical analysis**). Thus, left vTP lesioned patients showed a relative emotional amnesia despite a behaviourally derived index that emotional words were encountered as different from the others in the list.

### Emotional word amnesia is not explained by reduced volume in the amygdala or the hippocampus

The human temporal pole is strongly connected both structurally^40^ and functionally^41^ to medial temporal structures. Epileptic activity in the TP could, therefore, have led to dysfunction in amygdala and hippocampus, thereby contributing to emotional memory impairment. To test, at a structural level, whether emotional amnesia in the left vTP lesioned group could have been explained by a reduction in grey matter volume in the amygdala and hippocampus due to epilepsy or another possible consequence of left vTP lesions, we performed volumetry on pre-operative MRIs. There was no difference in total intracranial volume between the left vTP group (*n* = 5, including one patient with bilateral vTP lesions) and the control group (*n* = 26, Mann-Whitney *U* = 36, *P* = .129, *r* rank biserial (*rrb)* = −.45), so volumes of regions of interest were corrected by total intracranial volume prior to analysis. Importantly, the left vTP group did not differ from the control group in amygdala volume (left amygdala: *U* = 64, *P* = .979, *rrb =* −.02; right amygdala: *U* = 64, *P* = .979, *rrb* = −.02) or hippocampus volume (left hippocampus: *U* = 53, *P* = .548, *rrb* = −.18; right hippocampus: *U* = 57, *P* = .696, *rrb* = −.12). The focality of vTP lesions in our cohort is underscored by the absence of group differences in standard volumetric measures of temporal pole volume (left vTP: *U* = 90, *P* = .188, *rrb* = .38; right vTP: *U* = 82, *P* = .387, *rrb* = .26). However, the functional consequences of these small lesions may be more anatomically extensive (as shown by metabolic PET in patients r15 and r27 in **Supplementary Fig. 2**). Furthermore, in two out of the five patients with left vTP lesions, the lesions were caused by focal dysplasia, which is typically not related to reductions in gray matter volume. The grey matter volumes were also normal in the unilateral right vTP lesioned patient, with all structures being within 1 SD from the control group mean (left amygdala: *z* = −.43, right amygdala: *z* = .53, left hippocampus: *z* = .34, right hippocampus: *z* = .19).

### Left temporal pole lesions do not disrupt amygdala gamma responses to verbal stimuli

Having established structural integrity of amygdala and hippocampus in left vTP patients, we proceeded to measure whether functional responses, indexed by direct intracranial EEG (iEEG) recordings, in amygdala and hippocampus were similar in vTP lesioned and control patients or whether the reduction in verbal emotional memory could be explained by reduced functionality in these regions. Given the anterior temporal role in semantic processing, a critical question to address was: if the left vTP is damaged, will the amygdala “know” that a word is emotionally aversive. Previous research has related both passive viewing and successful encoding of verbal and visual negative memories to an increase in amygdala gamma power^6,42,43^. We first analysed time-resolved spectral responses to emotional words compared with control words in the amygdala during encoding in the verbal task.

Although there was no significant effect of emotion when including both the left and the right amygdala in this analysis, given the verbal nature of the task, and previous fMRI evidence for left unilateral amygdala responses to emotionally negative words^39^, we tested for a larger difference in gamma power between emotional and control words in the left hemisphere (*n* = 12) compared with the right hemisphere (*n* = 8), which we indeed observed around 450-980ms between 60-92.5Hz (cluster-permutation corrected *t*_sum_ = 1125.16, *P* = .028) (**Extended Data Fig. 3**). Post-hoc testing confirmed higher gamma power in the left amygdala for emotional compared with neutral control words between approximately 610-1080ms post stimulus presentation between 37.5-105Hz (*n* = 12, cluster-permutation corrected *t*_sum_ = 1061.72, *P* = .017; **Fig. 2g-h**). There was no significant difference between emotional words and perceptual oddballs in the left amygdala (all *p*s ≥ .794). In line with a left lateralized task, there was no effect of emotion in the right amygdala (*n* = 8, all *p*s ≥ .516), and an independent samples t-test showed that the difference in gamma power between emotional and neutral control words in the significant cluster was larger in the left (*n* = 12) than in the right (*n* = 8) hemisphere (*t*(18) = 2.86, *P* = .010, *d* = 1.30, **Fig. 2i**). We finally tested whether there was a difference in the emotional gamma effect between the left vTP group (*n* = 5) and the control group (*n* = 7) within the left amygdala (in the significant cluster from the emotional vs neutral control word analysis), and found no significant difference between the two groups (*U* = 28, *p* = .106, *rrb* = .60, **Fig. 2i**). These results again indicate that a relative emotional amnesia occurs in left vTP patients despite a physiologically-derived index at encoding (amygdala gamma activity) that emotional words were encountered as different from the others in the list, in line with slower reaction times for emotional items.

### Emotional visual episodic memory is unaffected by left temporal pole lesions

Having established an emotional memory impairment in a verbal memory task in patients with left vTP lesions, we next tested whether the memory impairment generalized to visual emotional memory. In this task, participants first viewed neutral and emotionally negative pictures while making indoor/outdoor judgements (**Fig. 3a**). Twenty-four hours later, the studied images were shown again, randomly intermixed with new images, in a remember/know recognition memory test to distinguish recognition based on recollection (remember), where details from the encoding episode are remembered, from recognition based on mere familiarity (know)^44^. Emotion selectively enhances recognition based on recollection^45^ and previous fMRI data has demonstrated that amygdala encoding-related responses are larger for subsequently recollected compared with familiar or forgotten stimuli^5^. Recognition memory scores were analysed using a Group (control, *n* = 19 vs left vTP, *n* = 4) × Emotion (negative vs neutral) × Response (remember vs know) mixed ANOVA (**Fig. 3b**). Importantly, there were no interactions involving group (all *P*s ≥ .304), indicating that, in contrast to verbal memory, there was no reduction in emotional memory in the visual domain in the left vTP lesioned group. There was a main effect of Emotion (*F*(1,21) = 7.04, *P* = .015, η_p_^2^ = .251), indicating that the proportion of hits was higher for aversive pictures compared with neutral pictures and a main effect of Response (*F*(1,21) = 7.19, *P* = .014, η_p_^2^ = .255) indicating that the proportion of hits was higher for remember than for know responses. When removing the factor of Group (which did not interact with any other factors from the ANOVA) we replicate the Emotion × Response interaction that we previously reported in a subset of this patient sample^6^ (*F*(1,22) = 4.40, *P* = .048, η_p_^2^ = .167. Follow-up pairwise comparisons showed that there was a difference in recognition memory between negative and neutral pictures for remember responses (t(22) = 3.06, *P* = .006, *d* = .64) while there was no such difference in recognition performance between negative and neutral pictures for know responses (*t*(22) = 1.00, *P* = .330, *d* = .21). Similarly to the accuracy results, there were no differences between the left vTP lesioned group and the control group in reaction times to visual stimuli during encoding or the recognition test (see **Extended Data Fig. 4**).

**Fig. 3.**
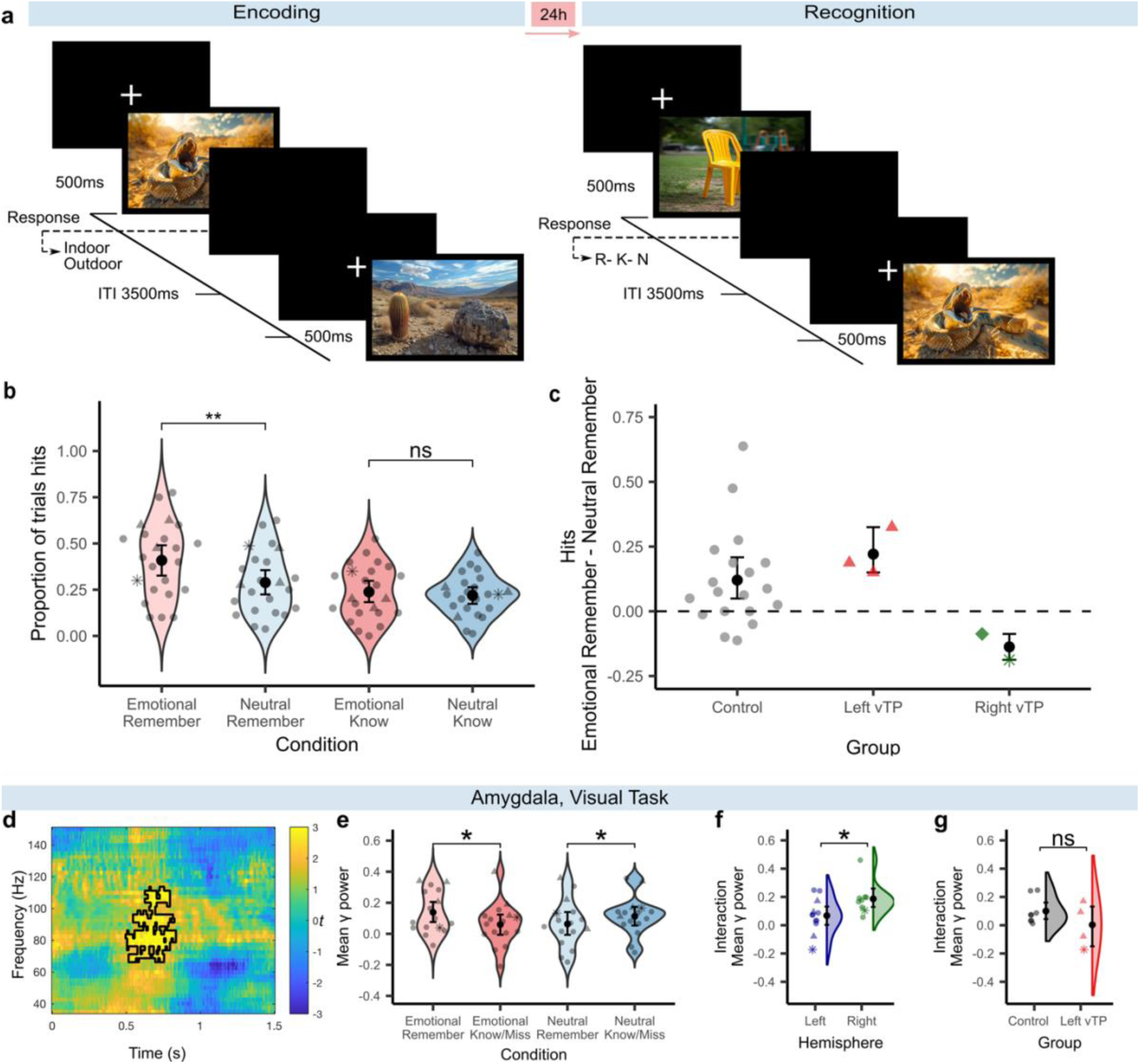
Experimental design, behavioural results and amygdala intracranial recordings in the visual emotional memory task. **a**, Illustration of the emotional memory task with emotional (negative) and neutral pictures. ITI = Inter Trial Interval **b**, Recognition memory performance, shown as the proportion of responses to old images that are hits for emotionally negative and neutral pictures as a function of response (remember/know). Scatter dots indicate individual means of proportion hits (▲ left vTP lesion, ♦ right vTP lesion, * bilateral TP lesions, • control patient, here and in subsequent panels). The black dots represent the group means of proportion hits and error bars depict the 95% confidence interval of the mean. Significance levels: *** < .001 and ns = not significant. **c,** Descriptive data of emotional memory performance in the control group, the left temporal pole group and the right temporal pole group. Note that the bilateral vTP patient is shown in the right vTP group in this figure. The y-axis depicts memory enhancement for negative versus neutral pictures for remembered responses (scores above 0 indicate emotional memory enhancement). **d,** Time-resolved spectral power induced by picture presentation (time 0ms) in bilateral amygdala. Colour scale indicates t-values from the comparison of emotionally negative vs neutral pictures, with black outline indicating a significant cluster from cluster-based permutation tests. **e**, Extracted mean gamma power from significant clusters. Black dots indicate group means and error bars represent 95% confidence interval of the mean. Significance levels: ** < .01, * < .05, ns = not significant. **f**, Mean gamma power from the significant cluster from the left and the right hemisphere. **g**, Mean gamma power from the significant cluster separated by group: control and left vTP (ventral temporal pole).

To summarize, the ANOVA results showed that visual emotional memory was equal in both groups, suggesting that the emotional memory impairment in left vTP patients was specific to the verbal domain. Intracranial electrophysiological analyses (described below) revealed a greater involvement of the amygdala in the right hemisphere in the visual episodic memory task compared with the left amygdala (**Fig. 3**). This led to a hypothesis that the two patients in our sample with right-sided vTP lesions (one with unilateral right and one with bilateral vTP lesions) would show impaired emotional memory in the visual task. Confirming our prediction, both these patients showed *reduced* memory for remember responses for emotional vs neutral pictures (**Fig. 3c)**. In contrast, only 4 of the 19 tested control patients (21.1%) and 0 of the 3 patients with unilateral left sided temporal pole lesions (0%) performing this visual task had lower remember performance for emotional compared with neutral images in this task. The reduction in emotional memory for remembered responses (emotional remembered minus neutral remembered) was −8.8% (z = −1.10 compared with the control group) in the patient with unilateral right temporal pole patient and −18.8% in the patient with bilateral temporal pole lesions (z = −1.63 compared with the control group). Taking this descriptive case analysis together with the inferential group analyses described above, suggest that the left vTP is involved in verbal emotional memory while the right vTP may be involved in visual emotional memory.

### Left temporal pole lesions do not disrupt amygdala gamma responses to visual stimuli

Bilateral amygdala gamma responses during the visual emotional memory task showed an Emotion (negative vs neutral) × Memory (remembered vs know/forgotten) interaction starting around 500ms and lasting until 830ms post stimulus presentation between 67-110Hz (*n* = 16, cluster-based permutation correction *t*_sum_ = 908.80, P = .033 cluster corrected; **Fig. 3d**) as previously reported in a smaller sample of the same patient cohort^6^ . Follow-up paired t-tests on mean power from the significant cluster showed that subsequent memory for negative pictures was related to increased gamma power (*t*(15) = 2.79, *P* = .014, *d* = .70, **Fig. 3e**) while for neutral pictures subsequent memory was related to reduced gamma power (*t*(15) = −2.84, *P* = .012, *d* = .71). In contrast to amygdala responses in the verbal task, this Emotion × Memory amygdala gamma effect in the visual task was larger in the right (*n* = 9) than in the left (*n* = 12) hemisphere (*t*(19) = 2.29, *P* = .034, *d* = 1.01, **Fig. 3f**). The left vTP group (*n* = 4) had contacts in the left amygdala, so we compared them to patients in the control group with contacts in the left hemisphere (*n* = 8). There was no difference in the size of the Emotion × Memory amygdala gamma interaction effect between the two groups (*U* = 20, *P* = .570, *rrb* = .25, **Fig. 3g**). The right amygdala gamma response in the patient with bilateral vTP lesions followed the same pattern as the rest of the patients (mean gamma power for interaction = .104, *z* compared to control group = −.79).

### Hippocampal gamma responses to verbal and visual stimuli

A subgroup of patients had also been implanted with electrodes in the hippocampus. Analyses of hippocampal iEEG data, during encoding, showed no difference between the left vTP lesioned group and the control group in theta responses related to saliency in the verbal task (both perceptual oddballs and emotional words vs neutral control words) and gamma responses to emotion in the visual task (**Extended Data Fig. 5**). There were also no group differences in hippocampal responses related to subsequent memory during encoding in both tasks (**Extended Data Fig. 6**). However, the number of left vTP lesioned patients with contacts in the hippocampus was too few to test deviance from the control group statistically in the visual task (two left vTP lesioned patients).

### A lateralized, modality-specific role for human vTP in emotional memory

Across verbal and visual memory tasks, memory is greater for emotionally negative vs neutral stimuli in patients without vTP lesions (control patients). Left vTP lesions result in worse memory for negative vs neutral words, but enhanced memory for negative pictures. A patient with a right vTP lesion shows the opposite pattern. A patient with bilateral vTP lesions shows worse memory for negative vs neutral words and pictures (**Fig. 4**).

**Fig. 4.**
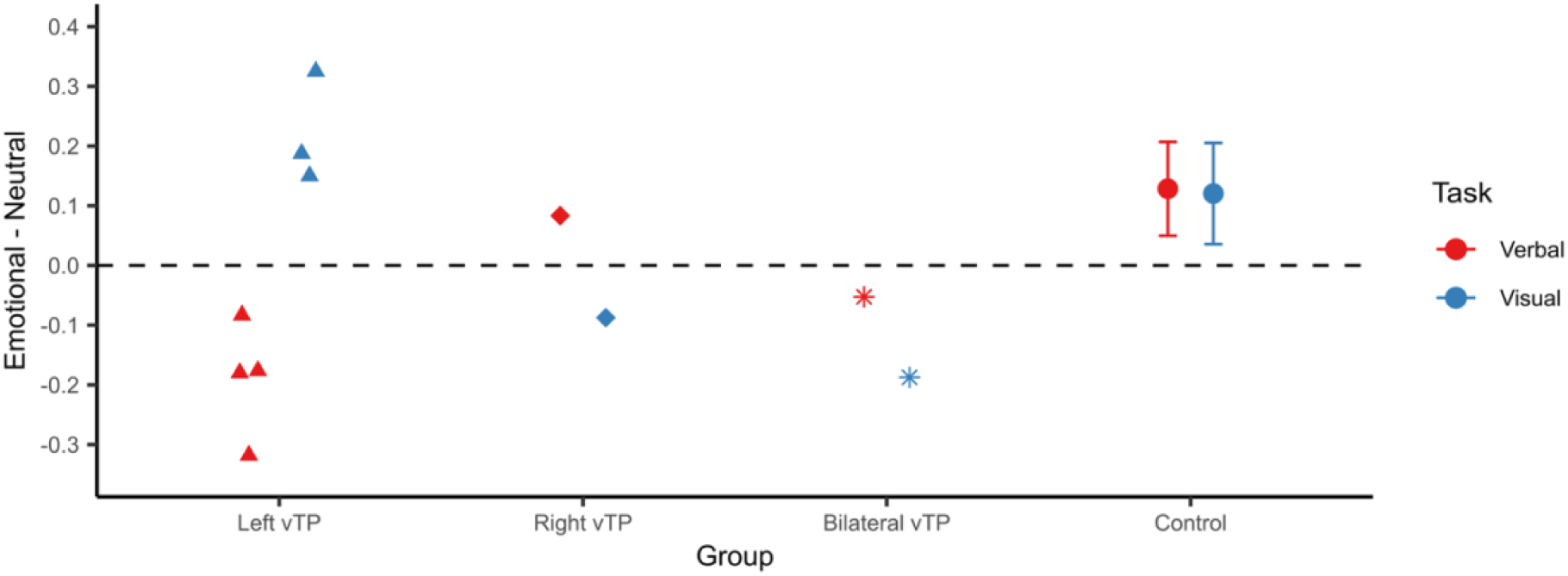
Summary of memory performance on verbal and visual memory tasks. Individual patient memory performance for emotional minus neutral items in the verbal (emotional minus neutral words) and the visual task (Remember responses to emotional images minus neutral images). ▲ left vTP group, ♦ right vTP group, * bilateral TP group, • control group. The mean and 95% confidence intervals are shown for the control group for each task.

## Discussion

The temporal pole is involved in emotional processes^10,17^, but little is known about its role in emotional memory. Determining the functional role of the TP is challenging when employing a neuropsychological approach, because pathologies affecting the TP typically also affect medio-temporal structures involved in emotional memory. Here, among a large, multi-site cohort of medication-resistant epilepsy patients performing emotional memory tasks, we identified 6 patients with vTP damage, 5 of whom had highly focal lesions. Patients with left vTP damage showed verbal emotional amnesia. This pattern of results, lower memory for emotional words than for neutral control words, was present in all 5 left vTP lesioned patients, but was absent in 1 patient with a unilateral right vTP lesion. By contrast, this patient with unilateral right vTP damage and 1 patient with bilateral vTP damage both showed lower memory for emotional vs neutral pictures in the visual memory task, whereas left unilateral vTP lesioned patients showed typical emotional picture memory enhancement (**Fig. 4**). These results are, therefore, suggestive of a double dissociation^46^ where the left vTP is involved in verbal emotional memory and the right vTP is involved in visual emotional memory.

Our understanding of how humans preferentially store emotional stimuli in long-term, episodic memory, derives largely from studies on single cases with focal, bilateral amygdala damage^2,3^, whose memory performance for arousing, emotional negative stimuli was reduced to the same level as that for neutral stimuli. Subsequent human fMRI and intracranial recording studies indicate an amygdala-dependent modulation of hippocampal activity underlying emotional memory^2,5,6,12^ . In order to propose a necessary and sufficient role for vTP in emotional memory it was therefore imperative to demonstrate absence of abnormality in amygdala and hippocampus in these patients (which could have been due to their epilepsy or as a downstream consequence of vTP damage). Inferring causality from brain lesions is generally limited by the alternative explanation of altered function of brain networks. In the current study, simultaneous intracranial recordings during emotional memory task performance confirmed that both amygdala and hippocampus of left vTP lesioned patients showed an increase in gamma responses during encoding of emotional words that was equivalent to control patients. Moreover, analysis of structural MRI data showed that the left vTP lesioned group had normal amygdala and hippocampus volumes. Taken together, the results indicate functionally and structurally intact amygdala and hippocampus in these patients thus their dysfunction is unlikely to explain the verbal emotional amnesia in the left vTP lesioned group. While the functional integrity of other brain structures potentially involved in emotional memory cannot be claimed, the current results go some way towards establishing the vTP as causally involved in episodic memory for emotionally negative stimuli.

The anterior temporal lobe, including the pole, is associated with semantic function, which is a broad term, with some models explicitly including emotion as a semantic feature dimension^47^ . An important observation here in this regard is that the impairment in verbal emotional memory in the left vTP group remained when controlling for other semantic word properties known to affect episodic memory including semantic association strength to the other words in the list, word frequency, word familiarity, concreteness and imageability. Memory performance was lower for the left vTP lesioned group than for the control group for 28 out of the 30 emotional words, indicating a general emotional memory impairment. Because word lists comprised semantically-related items, a semantic deficit in TP patients would predict worse memory for neutral words, due to either an inability to semantically link items at encoding, or a failure to benefit from a category-cued retrieval strategy. We show this was not the case: memory for neutral nouns was equivalent between left vTP and control groups. The patients with right vTP lesions showed impaired emotional, but preserved neutral, memory for pictures in the visual task, which did not involve semantically-related stimulus lists. Taking these observations together with the normal performance of left and right vTP patients in neuropsychological scoring on verbal comprehension, indicates that emotional memory deficits in vTP lesiones patients cannot be ascribed to a general semantic deficit. That memory for neutral words and pictures was spared in left and right TP patients, respectively, is also in keeping with a prevailing view that the temporal polar region does not play an important role in episodic memory for recent experiences^48^ (but see ^49–51^).

Our observations of a lateralized, modality-specific emotional memory impairment for verbal and visual stimuli following left and right vTP damage, respectively, informs ongoing debate regarding lateralization of left and right anterior temporal lobes in processing semantic memory from different sensory modalities or whether they process higher-level, amodal semantic representations. The amodal hub view is based primarily on the fact that patients with semantic dementia experience domain-general semantic impairments^16,17,52^ with bilateral atrophy causing greater impairment than unilateral atrophy^53^. Semantic dementia does, however, affect the anterior temporal lobes bilaterally, making it difficult to draw conclusions based on this patient group. Studies comparing semantic dementia patients with predominantly left or right atrophy have suggested that the left anterior temporal lobe is more involved in verbal processing while the right anterior temporal lobe is more involved in visual processing^54,55^. In addition, two recent larger studies have described that patients with predominantly right anterior temporal lobe atrophy show more emotion-related symptomatology (e.g. apathy, depression and lack of empathy) while patients with more pronounced left atrophy show more semantic symptoms such as anomia^19,20^. Our findings establish that both left and right anterior temporal lobes, and vTP specifically, are involved in emotional processing, in a modality-specific manner. Similar lateralization was evident in amygdala gamma responses during encoding of emotional items, with left amygdala responding more to emotional words than right amygdala, and *vice versa* for emotional pictures, so lateralization of vTP could be a manifestation of connectivity with other brain structures with lateralized processing^17^.

To arrive at a putative mechanistic role of vTP in emotional memory, it is important to note a key difference in effects of amygdala lesions from those described here. Amygdala lesions result in equal memory for neutral and emotional material^2,3^, whereas vTP lesion patients’ performance for emotional items was *lower* than for neutral items (i.e., emotional amnesia), suggesting that emotional stimuli have not simply lost a mnemonic benefit imbued by emotion. Thus, a different mechanism is needed to explain the emotional amnesia following vTP lesions. Here, we list evidence in favour of a possible failure of emotional memory retrieval. First, intracranial recordings indicate that the amygdala and the hippocampus responded normally (*i.e.,* not different from control patients) to emotional words (and pictures) during encoding. Second, reaction times were slower to emotional words during encoding suggesting that the left vTP lesioned patients still process these words as emotional during encoding. Drawing on evidence for TP involvement in affective/emotional semantics^17^ ^9^, it is possible that TP lesions lead to impaired retrieval of affective/emotional semantics. We propose that lack of access to category or semantic queues for emotional content due to damage in vTP leads to (near complete) failure of the episodic memory retrieval mechanism for emotional stimuli. This is in keeping with previous PET evidence for a left TP role in engaging in emotional retrieval mode^26^ and would explain how memory performance was reduced below the level of neutral items. We cannot measure iEEG activity during free recall of emotional words, or recollection-based “Remember” responses to emotional pictures in left and right vTP lesioned patients, respectively, as so few emotional items were retrieved.

To conclude, by studying focal vTP damage in a cohort of rare patients undergoing intracranial recordings, we present novel evidence for causal involvement of the vTP in emotional memory. Our observations highlight potential emotional memory deficits even in the earliest stages of neurological conditions affecting the TP and are relevant for presurgical evaluation in refractory epilepsy, as they predict specific impairments of emotional memory after resection of the vTP even if the amygdala and hippocampus are spared^56^. Furthermore, establishing a critical role of vTP in emotional memory has importance for the development of neurobiological models of mood disorders, including depression, which have been linked to reduced volume of the TP^8^. Lastly, our findings are relevant for the management of conditions such as PTSD, where exaggerated aversive memory retrieval can be maladaptive Focal neuromodulation of the amygdala is being developed as a novel treatment avenue for PTSD^57^. Given that emotional memory impairments following vTP lesions are even more pronounced than those ensuing from amygdala lesions, functionally inhibiting the vTP may prove effective in reducing unwanted emotionally negative memories.

## Methods

### Subjects

Thirty-two patients (16 females, mean age = 33.41, SD age = 12.37) with pharmaco-resistant epilepsy gave informed consent before participating in the study (see **Supplementary Table 1**). Patients were invited to participate from 3 clinical centres: the Ruber International Hospital in Madrid, Spain, the Swiss Epilepsy Centre in Zürich, Switzerland, and the Mara Hospital in Bielefeld, Germany. Four of these patients had left vTP lesions, one had a unilateral right vTP lesion and one patient had bilateral vTP lesions. The lesion in one patient in the left group was larger and extended to more posterior temporal inferiolateral regions, while all other patients in this group had focal lesions in the temporal pole. The unilateral right vTP patient was excluded from all statistical group comparisons except for the analyses specifically investigating the role of the right vTP.

The remaining 26 patients, without lesions in the temporal pole, were assigned to the control group (see **Supplementary Table 1** for a description of the location of the lesions for the patients in this group). There was no significant difference in age between the left temporal pole group (*M* = 30.4, *SD* = 17.74) and the control group (*M* = 33, *SD* = 10.54; *W* = 80, *p* = .435). The patients had normal or corrected-to-normal vision and all patients except one in the control group were right-handed. Electrode implantation sites were chosen solely based on clinical criteria. The participants did not receive financial compensation for participating. The study had full approval from the local ethics committees of the Hospital Ruber Internacional, Madrid, Spain, the Kantonale Ethikkommission, Zürich, Switzerland (PB-2016-02055) and Bielefeld University Ethics Committee (EUB-2021-29) with oversight from the European Research Council Ethics Board. Patient b1 performed the verbal memory task whilst aged 15 years and repeated the consent procedure after turning 18.

Five left vTP patients, one right vTP patient and 16 control patients participated in the verbal task. Amygdala iEEG activity was recorded from 12 control patients and 5 left vTP patients, while hippocampus iEEG activity was recorded from 10 control patients and 4 left vTP patients.

Four left vTP patients, one right vTP patient and 19 control patients participated in the visual task. Amygdala iEEG was recorded in 4 left vTP patients and 12 control patients while hippocampus iEEG was recorded from 2 left vTP patients and 12 control patients in this task. See **Supplementary Tables 1-7** for details regarding which tasks were performed by each patient and whether they had amygdala and/or hippocampal iEEG recordings.

### Neuropsychological assessment

Patients underwent neuropsychological testing prior to electrode implantation, with intelligence and verbal comprehension assessed employing Wechsler Intelligence Tests. The participants were tested in their native language (Spanish or German). The verbal comprehension index was computed based on the information, vocabulary, comprehension and similarities subtests. Both the full-scale IQ and the verbal comprehension scores are standardized (on a scale with mean = 100 and standard deviation = 15) and controlled for age. Different versions of the Wechsler Intelligence test were used. Eleven patients were tested with WAIS-III (Wechsler Adult Intelligence Scale), 16 patients were tested with WAIS-IV, 1 patient was tested with HAWI (Hamburg Wechsler Intelligence test for adults), 2 patients were tested with WIE (Wechsler Intelligence test for adults) and 1 patient was under 18 years old and was tested with WISC-V (Wechsler Intelligence Scale for Children). All vTP lesioned patients had both full scale IQ and verbal comprehension scores. Three control patients were missing full scale IQ scores and were excluded from the analysis (final *n* = 23 control patients) and five control subjects were excluded from the verbal comprehension analysis due to missing data (final *n* = 21 control patients).

### Surgical procedure

At the Ruber International Hospital in Madrid, Spain, a contrast enhanced MRI was performed pre-operatively under stereotactic conditions to map vascular structures ahead of electrode implantation and to calculate stereotactic coordinates for trajectories using the Neuroplan system (Integra Radionics). Dixi Medical Micro-deep depth electrodes (multi-contact, semi rigid with a diameter of 0.8mm, contact length of 2mm, inter-contact isolator length 1.5mm) were implanted based on the Leksell method. At the Swiss Epilepsy Centre in Zürich, Switzerland, depth electrodes (1.3mm diameter, 8 contacts of 1.6mm length and with 3mm spacing between the two most distal contact centres and 5mm between all others) were stereotactically implanted. Finally, at the Mara Hospital in Bielefeld, Germany, Ad-Tech depth electrodes (1.1 mm diameter, 2.4mm contact length, 10-11 contacts per electrode and 4.5mm spacing between contact centres) were stereotactically implanted into the amygdala and the hippocampus.

### Electrode contact localization

The pre-operative T1-weighted magnetic resonance images (pre-MRI) were co-registered with the post-operative computer tomography (post-CT) images. Both the pre-MRI and post-CT images were skull-stripped by filtering out all voxels with signal intensities between 100 and 1300 HU and by spatially normalizing the image to MNI space using the New Segment algorithm in SPM8. Next, the resultant inverse normalization parameters were implemented to the brain mask supplied in SPM8 and the mask was converted into native space. We filtered out all voxels outside the brain mask and with a signal value in the highest 15^th^ percentile in the pre-MRI. The skull-stripped pre-MRI was then co-registered and re-sliced to the skull-stripped post-CT space. Next, we thresholded the post-CT to only display electrode contacts and overlaid the two images.

### Electrode contact visualization

We visualized the contacts using Lead-DBS^58^. To reconstruct and visualize the electrodes we selected the electrode model (Dixi or Ad-Tech), co-registered CT to MRI using the advanced normalization tools (ANTs) and subsequently the volumes were normalized to MNI ICBM 2009b nonlinear asymmetrical space based on preoperative MRI data. Brain shift correction was performed using Lead-DBS. We initially pre-reconstructed the electrodes using semi-automatic reconstruction. The reconstructed electrodes were inspected visually, and any misalignments were adjusted manually. This adjustment was based on postoperative CT, ensuring the trajectory was placed as accurately as possible within the centre of the electrode artifact. To visualize all the electrodes, we employed the Lead-group software. After this procedure, MNI coordinates were assigned to each electrode contact. In the final step, we plotted only the contacts for each electrode within the regions of interest, either the amygdala or the hippocampus.

### Intracranial EEG data acquisition

At the Ruber international hospital, Madrid, continuous intracranial EEG was acquired using an XLTEK EMU128FS amplifier (XLTEK, Oakville, Ontario, Canada). The iEEG data was recorded at a sampling rate of 500Hz from each electrode contact site with an online bandpass filter between 0.1-150Hz. If a patient was recorded with higher sampling rate, the data were down-sampled to 500Hz. Data recorded at the Swiss Epilepsy Center in Zürich were recorded with a Neuralynx Atlas system with a sampling rate of 4000Hz against a common average intracranial reference using an online band pass filter of 0.5-1000Hz. The data were later down-sampled to 500Hz. At the Mara hospital in Bielefeld, the data were recorded at a sampling rate of 2000Hz using a Compumedics Neuvo amplifier with no online filter.

### Pre-processing of intracranial EEG data

The preprocessing and analysis of intracranial EEG data was performed using the FieldTrip toolbox^59^ using MATLAB (version, 2024b). The iEEG were transformed to bipolar channels by subtracting signals from adjacent electrode contacts within the same structure (the amygdala or the hippocampus). Prior studies have shown that bipolar referencing optimizes estimates of local activity^60–62^. Epochs were extracted from −6 seconds to 7 seconds post stimulus presentation in the verbal task and from −7.5 seconds to 7.5 seconds post stimulus presentation in the visual task. The data were then detrended. No off-line filtering was applied. Trials containing artifacts, such as epileptic spikes, were detected and removed based on visual inspection in the time domain.

### Time-frequency decomposition

For low frequencies (1-34Hz) we used a sliding fast Fourier transformation with a Hanning taper as implemented in the mtmconvol function in FieldTrip^59^. The decomposition was performed between 1-34Hz in steps of 1Hz and the length of the sliding time window was 7 cycles. In contrast the higher frequencies (35-150Hz) were decomposed using 7 Slepian multi-tapers with a sliding time window of 400ms and a 10Hz frequency smoothing. Both the low and the high frequencies were decomposed in a time window between 1500ms pre stimulus presentation to 3000ms post stimulus presentation in steps of 10ms. The time-frequency data were baseline corrected by calculating the relative percentage change compared with the baseline period between −1000ms to −100ms prior to stimulus onset. The time-frequency data was visually inspected and trials with artefacts were removed (e.g. epileptiform activity and broadband noise from hospital equipment). The time-frequency data was then averaged over channels within each brain structure (separately for the amygdala and the hippocampus) and over trials within each condition.

### Stimuli

#### Words presented during the verbal emotional memory task

The word stimuli were originally selected from the Edinburgh associative thesaurus^2^ and were later translated to Spanish and used in one prior study^63^. The Spanish word stimuli were rated for valence by an independent sample of 19 native Spanish speakers (13 females, mean age = 32.3, SD age = 6.1) using the self-assessment manikin^64^ on a scale between one (negative valence) to nine (positive valence). The average valence rating with s.e.m given in parentheses) was 6.59 (.06) for neutral words and 1.83 (.11) for emotional words. The same participants also rated the semantic association strength between words within each list. One list of 14 words were shown at a time and the participants were asked to first read through all words and then rate how strong the semantic association was between each word and the general context of the list between 0 (no relation) to 10 (very strong relation).

Word frequency per million words was obtained for all 420 words from the EsPal database^37^. The log10 word frequency was used in the GLMM analysis to facilitate convergence. Subjective ratings of word familiarity, imageability and concreteness were obtained from the same database. These ratings were available for 322 of the 420 words (76.7%).

Patient b1 was tested with a German version of this task^2,65^ as this patient was a native German speaker. A cosine semantic similarity measure was derived from the German version of the FastText model^66^. The average similarity to the other words presented in the same list was computed for each word. Word frequency per million tokens was obtained from the dlexDB lexical database^67^, which includes sources from fiction, newspapers, scientific publications and functional literature. The log10 of the word frequency per million tokens was used in the analysis. A measure of word familiarity was obtained from the same database based on the cumulative frequency of all words of the same length sharing the same initial trigram. Subjective ratings of concreteness and imageability were obtained from the Automatically Generated Norms^68^. We conducted a control analysis both with and without the German patient (b1) given that the German word measures where not fully corresponding to the Spanish word measures (e.g. subjective ratings of semantic similarity and familiarity for the Spanish words and corpus driven objective measures for the German words). All measures were z-transformed on a single-patient level to put the Spanish and German measures on the same scale prior to the analysis.

#### Pictures presented during the visual emotional memory task

The stimuli consisted of 80 negatively valanced highly-arousing aversive pictures and 160 neutral low-arousing pictures. All stimuli except for eleven neutral pictures (landscape pictures selected from the internet) were selected from the International Affective Picture System (IAPS)^69^. Mean normative IAPS picture valence ratings were 2.04 (.05) for aversive pictures and 5.05 (.05) for neutral pictures. The mean arousal rating was 3.29 (.06) for aversive pictures and 6.3 for neutral pictures.

### Procedure

The stimuli were presented on a 27 × 20.3 cm monitor with a 1024 × 768 resolution at a distance of 50 cm from the patient’s eyes (30.2° – 22.9°). The patients were asked to fixate their gaze in the centre of the screen and to remain as still as possible and to avoid verbalizations during the encoding phases of both tasks.

### Verbal emotional free recall task

This task was performed by patients at the Ruber International Hospital in Madrid, Spain, and the Mara Hospital in Bielefeld, Germany. The participants were presented with 14 nouns, shown one at the time centrally on the screen in black font on a white background. The words were displayed for 1000ms followed by a blank screen for 2000ms. In a shallow encoding task, the participants were instructed to indicate via button presses whether the initial letter of each word contained an enclosed space. The participants were also told that their memory would be tested for the words and that they should not use any mnemonic strategy except for repeating the words when encoding them. This encoding phase of each list was followed by the presentation of a number, and participants were asked to count backwards out loud in steps of three during 30s. Immediately after this distractor task, instructions appeared on-screen to say as many words as could be remembered in a free recall task while the experimenter noted their responses. The free recall test was self-paced. This procedure was repeated for up to 30 lists. Some patients did not complete the full experiment. The number of completed lists per patient is presented in the **Supplementary Table 1**.

All 14 words in each list belonged to the same semantic category, but each of the 30 lists contained words from different semantic categories. In every list, twelve words were emotionally neutral and presented in the Times New Roman font. These words are referred to as standard words. In addition, each list included one emotional word, a negatively valanced emotional word presented in the same font. A perceptual oddball, a word presented in a different font was also presented in each list. There were 15 different fonts for the perceptual oddballs which were repeated in the second half of the experiment. Out of the 12 standard words in each list, two were selected as control words used in the analyses. Like the oddballs, these control words could not occur within the first five words in a list to control for primacy effects. Moreover, like oddballs they could not be presented immediately before or after an oddball, to control for potential carryover effects of oddballs.

### Visual emotional recognition task

This task was performed by patients at the Ruber International Hospital in Madrid, Spain, and the Swiss Epilepsy Centre in Zürich, Switzerland. Behavioural results from partly overlapping patient cohorts from this task have been reported in two previous publications from our group^6,70^. This task was divided into one encoding and one retrieval phase. In the encoding phase, each trial started with a fixation cross shown in the centre of the screen in white on a black background. Next, a negatively valenced or a neutral picture was shown on-screen for 500ms followed by a blank black screen for 3500ms. The patients were instructed to indicate via button presses if the displayed picture was an indoor or an outdoor scene. After 24 hours, the patients were tested using a remember/know task. They were shown all the pictures from the encoding phase (40 emotional and 80 neutral pictures) together with as many new foils (40 emotional and 80 neutral), randomly intermixed, and were instructed to respond *remember if* they remember seeing the picture the previous day, or to respond *know* if a picture seemed familiar although they could not remember seeing it or to respond *new* if they had not seen the picture during the encoding phase.

### Data analysis

All statistical tests were two-sided.

#### Behavioural data analysis

Memory performance in the verbal task was analysed using a binomial generalized linear mixed effect model using the lme4 package^71^ in R (version 4.3.1, R Core Team, https://www.r-project.org/). Memory performance was used as the dependent variable and there were two fixed effects of Group (left temporal pole group vs control group) and Condition (emotional oddball vs perceptual oddball vs control) and a random intercept effect of Patient ID. The model formula was:

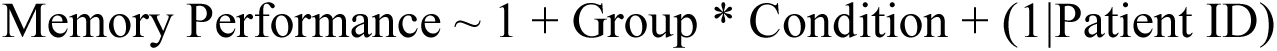

The statistical significance of the fixed effects and their interactions was assessed with an omnibus χ^2^ Wald test using the car package^72^. Significant interactions for both general mixed effects models and the mixed ANOVA analysis described below were followed-up using pairwise contrasts as implemented in the emmeans package^73^ applying a Benjamini & Hochberg (false discovery rate, FDR) correction for multiple comparisons.

Two additional GLMM analyses of memory performance in the verbal task were conducted to control for variables known to affect episodic memory. In the first analysis semantic association strength to the words in the same list and word frequency were added as co-variates. In the second analysis additional co-variates of word familiarity, imageability and concreteness were added as co-variates. The reason for conducting these two analyses separately was that the subjective ratings included in the second analysis were not available for all words. Bobyqa optimizer was used in both these GLMM analyses with co-variates.

The reaction times in the verbal free recall task deviated from a normal distribution as indicated by a Shapiro-Wilk test (W = .92, *P* < .001). The data were positively skewed and were analysed using a generalized linear mixed effect model with a gamma distribution and a log-link function. Incorrect trials and trials with a reaction time more than ± 3 SD from the mean were excluded from the analysis. The model included Group (left temporal pole group vs control group) and Condition (emotional oddball vs perceptual oddball vs control) and a random intercept effect of Patient ID:

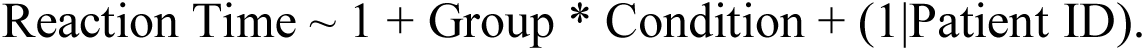

Memory performance in the visual recognition task was analysed using a mixed ANOVA. As previously described in Costa et al. 2022^6^ in a subsample of this data, there were significantly more false alarms for aversive compared with neutral pictures (*t*(22) = 4.78, *P* < .001, d = .996). One potential reason for the increase in false alarms for emotional items is that emotional items may be perceived as more related compared with neutral items, which may lead to an increased false sense of recollection due to overgeneralization^74,75^. It has however been shown that relatedness alone cannot explain all emotion-dependent differences in false recognition^76,77^. We analysed memory performance using a mixed ANOVA with factors of Group (left temporal pole vs control) × Emotion (aversive vs neutral) × Response (*remember* vs *know*). The false alarm analysis is presented in **Extended Data** Fig. 6.

#### iEEG data analysis

The sample included in each analysis varies between different structures, as all patients did not have electrodes in both the amygdala and in the hippocampus, and varies between the left and the right hemisphere as most patients only had electrodes in one hemisphere. The data were averaged for all bipolar channels within a structure in each patient. In patients with bilateral recordings, the left and the right channel were averaged for each structure in the initial analyses, but analysed separately when comparing the left and the right hemisphere and when comparing the control group to the left vTP group (given that all but one left vTP patient only had left sided recordings). All iEEG data were recorded during the encoding phase in both tasks.

In the verbal task, we analysed electophysiological activity from the amygdala in 12 control subjects (2 bilateral, 5 left sided and 5 right sided) and 5 left vTP patients (1 bilateral and 4 left sided) and from the hippocampus in 10 control patients (1 bilateral, 3 left sided and 6 right sided) and 4 left vTP patients (all with electrodes in the left hippocampus). In the visual task, we analysed iEEG activity from the amygdala from 12 control patients (4 bilateral, 4 left sided and 4 right sided) and 4 left vTP patients (1 bilateral, 3 left sided) and from the hippocampus in 12 control patients (2 bilateral, 4 left sided, 6 right sided) and 2 vTP patients (both left sided). See **Supplementary Tables 1-7** for detailed descriptions of which patients were included in the iEEG analyses of the amygdala and the hippocampus in each task.

Our planned comparisons in the verbal task were between emotional nouns and the control words. The iEEG time-frequency analysis was conducted using cluster-based permutation tests as implemented in FieldTrip^78^. Lower frequencies (1-34Hz) and higher frequencies (35-150Hz) were analysed separately within each structure (the amygdala and the hippocampus). First, t-tests were conducted at time/frequency datapoint. Neighbouring samples in time and/or frequency were combined into clusters and the sum of all samples t-values was computed. The observed cluster-level t-value was next compared with the t-value in 10000 permutations (or the maximum number of unique permutations if lower than this value) and the Monte Carlo *p*-value was computed as the proportion of permuted cluster-level t-values being larger than the observed cluster-level t-value. Cluster-based permutation tests do not provide statistical inference for the exact latency and frequency of the effects^79^. All tests were two tailed and the p-values are reported with alpha = 0.05. All analyses followed the following logic: If there was an effect when averaging over both left and right recordings, then the power values were extracted from the significant cluster and then we assessed whether the effect was larger in the left compared with the right hemisphere and in the left vTP group compared with the control group using independent t-tests. A Mann-Whitney U test was used instead of an independent samples t-test when the assumption of normality was violated as indicated by a Shapiro-Wilk test of normality and when there were large differences in group sizes (when the control group was more than twice as large as the vTP group). If there was no bilateral effect, as in the analysis of the emotion effect in the amygdala, then we performed a cluster permutation test to see if the effect was larger in one of the hemispheres (the left in the verbal task). Next, we ran a cluster-permutation test within the hemisphere where the effect was the largest (the left hemisphere) and extracted the power values from this cluster for testing if there was any difference in the effect between the left vTP group and the control group.

A similar approach was used in the visual task. As described in Costa et al. 2022^6^ we had an Emotion (aversive vs neutral) × Subsequent memory (remembered vs forgotten) design and analysed the encoding data with cluster-based permutation tests. Given the low number of *know* responses for emotional items, know items were collapsed with forgotten trials for both aversive and neutral pictures. Significant interactions were followed-up by extracting the power from the significant cluster and comparing the conditions with paired t-tests. Next, we tested if the interaction effect was lateralized with a t-test on the difference between remembered and forgotten items subtracted between aversive and neutral pictures. Finally, we investigated if the interaction was larger for left temporal pole subjects than for control patients (analysing only contacts from the left hemisphere to be comparable with the left temporal pole patients). We also calculated the *z*-score for the right amygdala response in the patient with bilateral vTP lesions given that this patient had reduced visual emotional memory. The unilateral right vTP lesioned patient could not be included in this analysis, due to insufficient signal quality in the right amygdala iEEG data.

### Regional volume analyses

Volumes of regions of interest (amygdala, hippocampus and temporal pole cortical grey matter, left and right in all cases) were obtained using FreeSurfer^80^ 7.1.1. Pre-operative T1-weighted MRI scans from each patient were processed using the standard *recon-all* FreeSurfer pipeline, which provides cortical and subcortical segmentations. This was followed by specific sub-segmentation of the amygdala^81^ and hippocampus^82^ to retrieve more accurate volumes for these two structures. Volumes (in cubic mm) were then obtained from these segmentations and corrected as a function of total intracranial volume. Specifically, regional volumes were regressed by intracranial volume in a linear model, and corrected volumes were calculated as original volumes plus model residuals. Corrected volumes per region were then compared among groups using the Mann-Whitney U test. All patients except the patient with unilateral right vTP lesion were included in the analysis.

### Lesion mapping

The spatial distributions of focal dysplasia, encephalocele and traumatic brain injury were manually segmented using T1 MRI and FDG-PET images employing MRIcron^83^. The T1-weighted MR data, and their respective lesion segmentations, were then resampled to the same voxel dimensions as the MNI152 template, before the T1-weighted MR image underwent brain extraction using the HD-BET^84^ preprocessing method. This extracted T1 brain image was then registered to MNI co-ordinate space using the SyNOnly (Symmetric normalization with no rigid or affine stages) method of the Advanced Normalization Techniques Python library (ANTsPy)^85^, having computed the image registration across the range of available methods and selecting SyNOnly based on visual inspection. The transformation fitted using the T1 image was then applied to each respective lesion segmentation image.

The Brodmann atlas, packaged with MRIcron^83^, already in MNI co-ordinate space, was used to extract the Brodmann Areas (BAs) affected by the registered lesions. The proportion of the volume of each lesion, located inside each BA was recorded. Where the lesion protruded outside the defined BA regions, the nearest defined voxel, by Euclidean distance, was instead recorded. This therefore facilitated a decomposition of each lesion by the affected BAs, according to the respective proportion of the lesion volume. The same procedure was used to estimate the affected AICHA areas^31^. The overlap between lesions and BAs and AICHAs are shown in **Extended Data Fig. 1 and 2**, respectively.

Lesion visualization was performed using MRIcroGL^86^ 1.2.2 and with ParaView^87^ 5.13.3, following conversion of the registered lesions from NIfTI to VTK using ITK-SNAP^88^ 4.0.1.

## Data availability

The electrophysiological data generated in this study will be deposited at the project’s open science framework page when the manuscript is accepted for publication in a scientific journal. The raw patient data are protected and not available due to data privacy regulations and the conditions of ethical approval. Additional processed data supporting the findings of this study will be available at the same link. Further information can be obtained by contacting the lead author.

## Code availability

The MATLAB and R scripts used to analyze the data in this study will be publicly available on the project’s open science framework page when the manuscript is accepted for publication in a scientific journal.

## Acknowledgements

This project has received funding from the European Research Council (ERC) under the European Union’s Horizon 2020 research and innovation programme (ERC-2018-COG 819814) to BAS. JS was funded by the Swiss National Science Foundation (SNSF 204651). The example pictures in **Fig. 2a** were downloaded from www.stockcake.com. We thank all the patients for their participation.

## Author contributions

B.A.S designed the experiments and analyses, and acquired the funding. J.S., D.L, M.C., A.P.-Y., R.H., P.G., J.K. and R.T. collected data. A.G.-N. R.T., L.I., F.G.W and C.G.B. monitored patients and performed clinical evaluation. R.H. preprocessed the iEEG data from the verbal task and M.C. preprocessed the iEEG data from the visual task. D.O-C. processed the structural MRI data. R.H. manually segmented and D.G. normalised the temporal pole lesions. R.H. analysed and visualised the data. R.H. and B.A.S wrote the manuscript. All of the authors provided comments on the manuscript.

Supplementary Information is available for this paper.

## Competing interests

The authors declare no competing interests.

Correspondence and requests for materials should be addressed to Robin Hellerstedt or Bryan Strange.

## Notes

### Competing Interest Statement

The authors have declared no competing interest.

