## Extended Data for "Emotional amnesia in humans with focal temporal pole lesions"

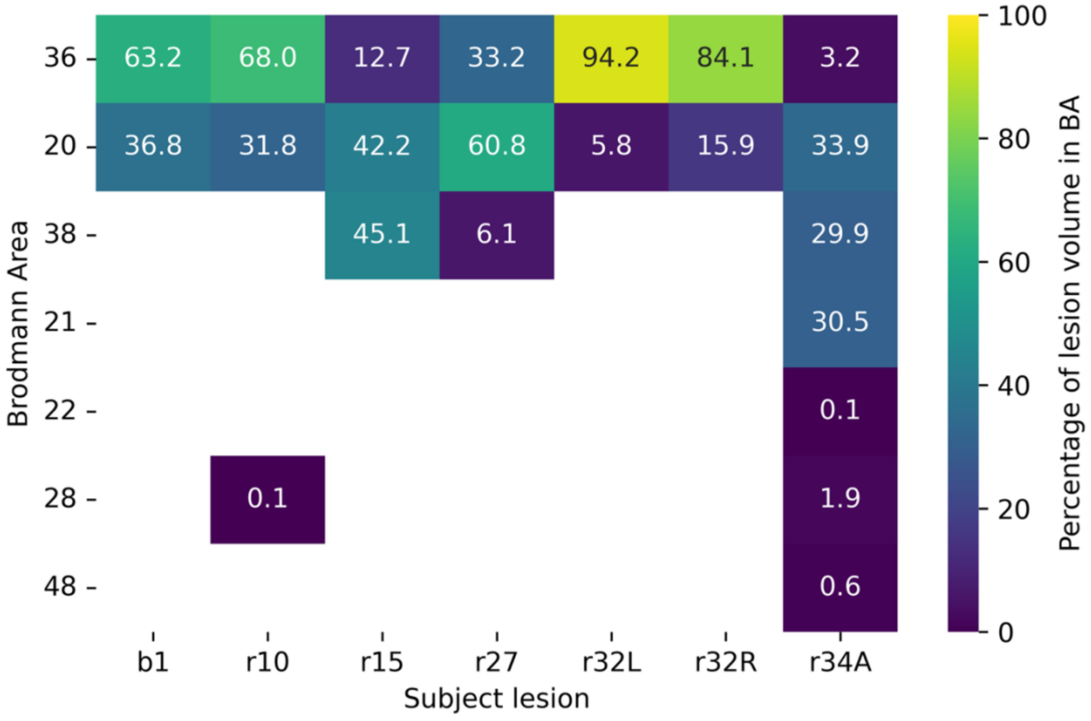


**Extended Data Fig. 1 | Overlap between lesions and Brodmann Areas.** The percentage of overlap between the lesion and Brodmann areas is shown for each lesion. Abbreviations in patient IDs: b = Mara Hospital in Bielefeld, r = Ruber International Hospital in Madrid, L = left hemisphere, R = right hemisphere, A = anterior.

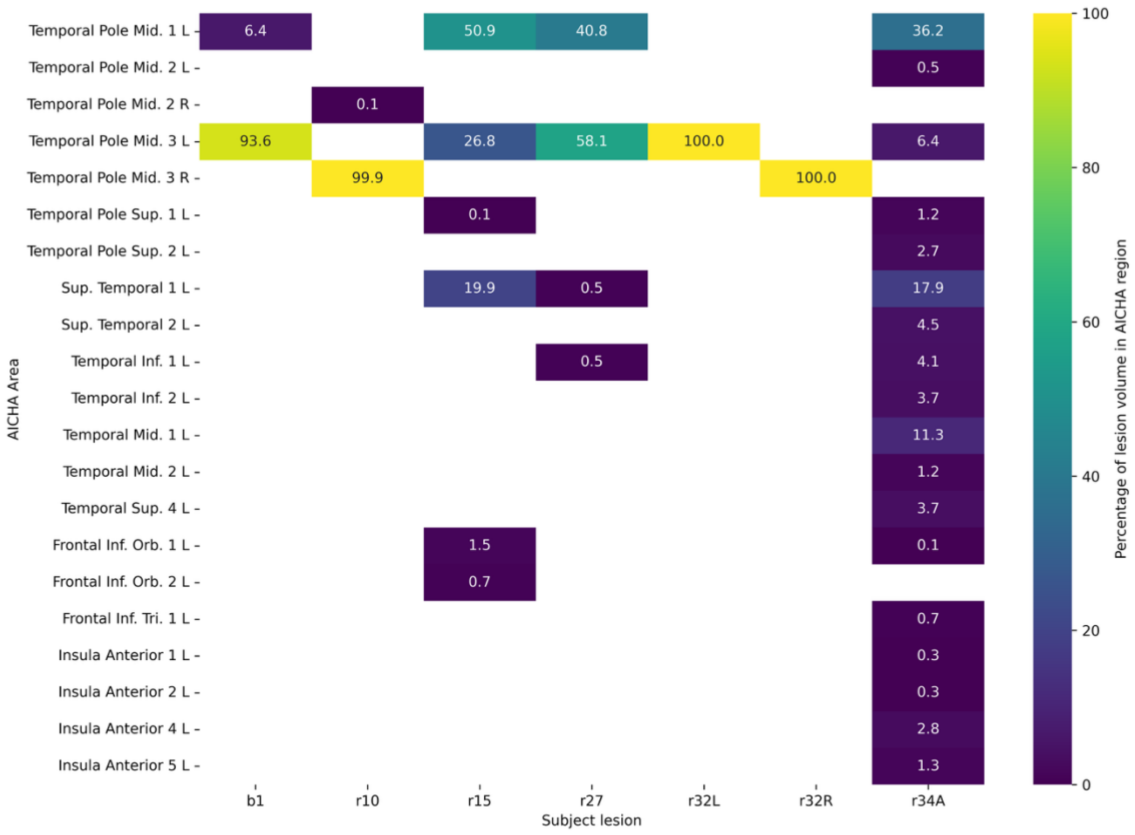


**Extended Data Fig. 2 | Overlap between lesions and regions in the Atlas of Intrinsic Connectivity of Homotropic Areas (AICHA).** The percentage of overlap between the lesion and AICHA areas is shown for each lesion. Abbreviations in patient IDs: b = Mara Hospital in Bielefeld, r = Ruber International Hospital in Madrid, L = left hemisphere, R = right hemisphere, A = anterior.

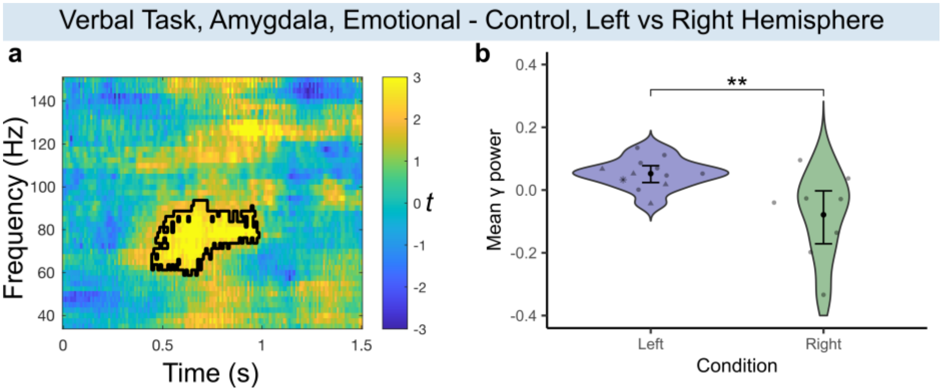


**Extended Data Fig. 3 | Left vs right amygdala responses to emotional words. a)** Time frequency plots showing t-values from cluster-based permutation tests between left hemisphere (emotional-control) and right hemisphere (emotional-control). Black outline indicates a significant cluster ($t_{sum}$ = 1125.16, $P$ = .028) around 450-980ms between 60-92.5Hz. **b)** Extracted mean gamma power from the significant cluster. Scatter dots display individual means (• control patient, ▲ unilateral left vTP lesion, * bilateral vTP lesion), black dots indicate group means and the error bars represent 95% confidence interval of the mean. Significance level: ** < .01.

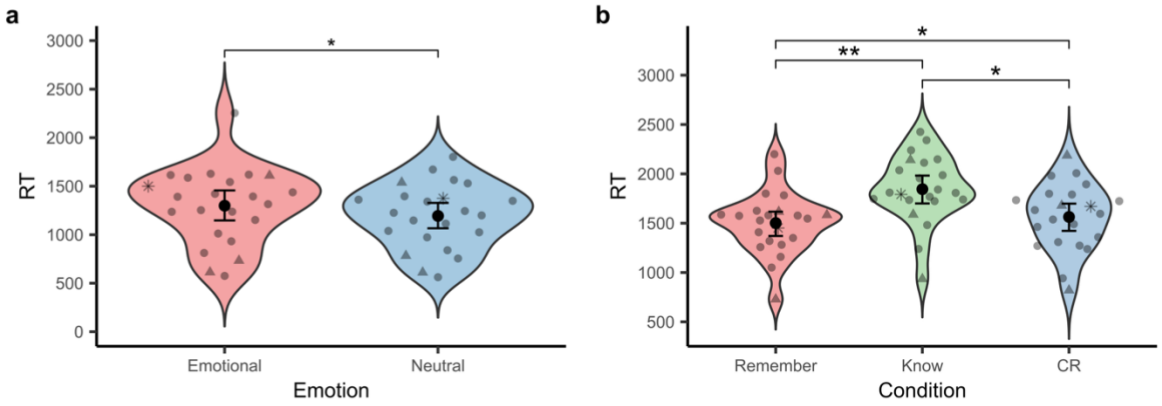


**Extended Data Fig. 4 | Reaction times from the visual task. a)** Reaction times from the indoor/outdoor task during encoding. Scatter dots display individual means (• control patient, ▲ unilateral left vTP lesion, * bilateral vTP lesion), black dots indicate group means and the error bars represent 95% confidence interval of the mean. Significance levels: * < .05. **b)** Reaction times from the recognition test in the visual task. Significance levels: ** < .01, * < .05. The data analyses of reaction times from the visual task are described in **Supplementary Data 2**.

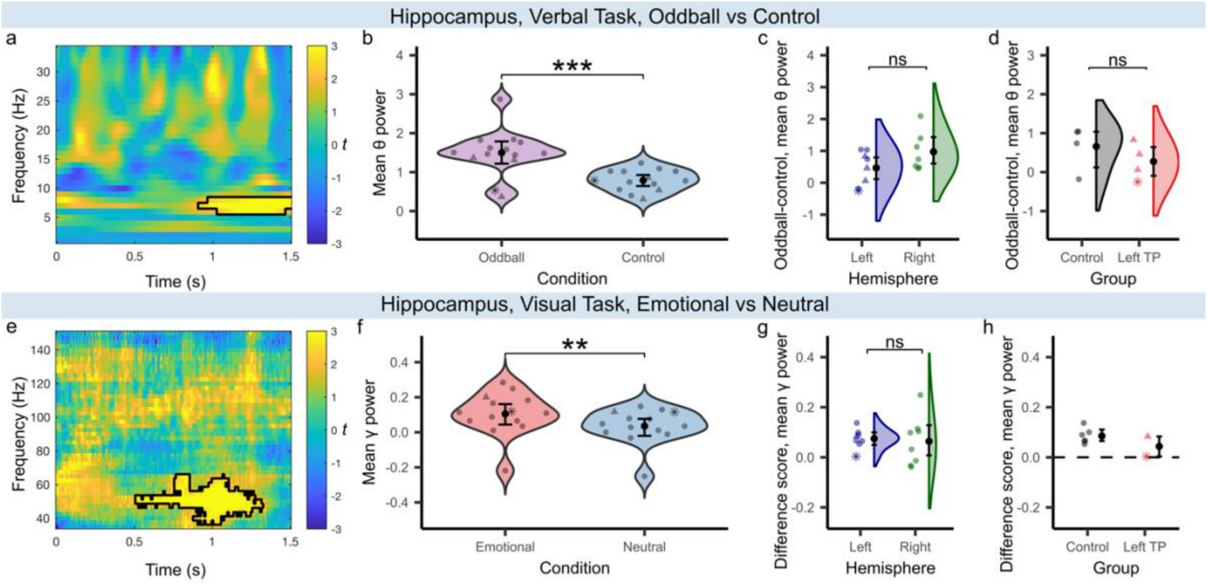


**Extended Data Fig. 5 | Hippocampus responses to saliency and emotion.** column 1: **a, e,** Time frequency plots showing t-values from cluster-based permutation tests. Black outline indicates a significant cluster. column 2: **b, f,** Extracted mean gamma power from significant clusters. Scatter dots display individual means (• control patient, ▲ unilateral left vTP lesion, * bilateral vTP lesion), black dots indicate group means and the error bars represent 95% confidence interval of the mean. Significance levels: *** < .001, ** < .01, ns = not significant. column 3**: c,g,** Mean gamma power from the significant cluster from the left and the right hemisphere. column 4: **d,h**, Mean gamma power from the significant cluster separated by group (control and Left vTP (ventral temporal pole)). The analyses of hippocampal responses to saliency and emotion are described in **Supplementary Data 3**.

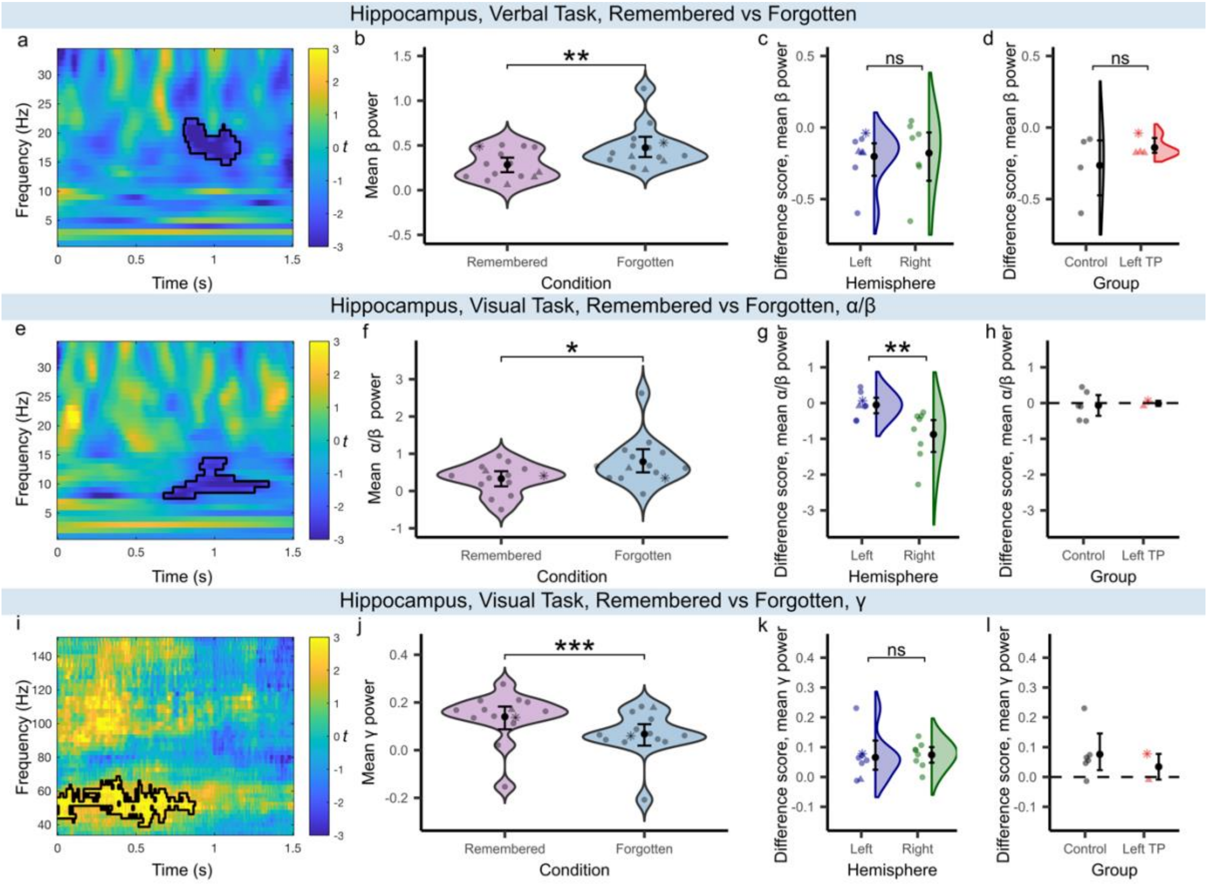


**Extended Data Fig. 6 | Hippocampus responses to subsequently remembered stimuli are not affected by left temporal pole lesions.** Column 1: **a, e, i,** Time frequency plots showing t-values from cluster-based permutation tests. Black outline indicates a significant cluster. column 2: **b, f, j**, Extracted mean gamma power from significant clusters. Scatter dots display individual means (• control patient, ▲ unilateral left vTP lesion, * bilateral vTP lesion), black dots indicate group means and the error bars represent 95% confidence interval of the mean. Significance levels: *** < .001, ** < .01, * < .05, ns = not significant. column 3**: c,g,k,** Mean gamma power from the significant cluster from the left and the right hemisphere. column 4: **d,h,l,** Mean gamma power from the significant cluster separated by group; control and left ventral temporal pole (vTP). The analyses of hippocampal responses to subsequent memory are described in **Supplementary Data 3**.

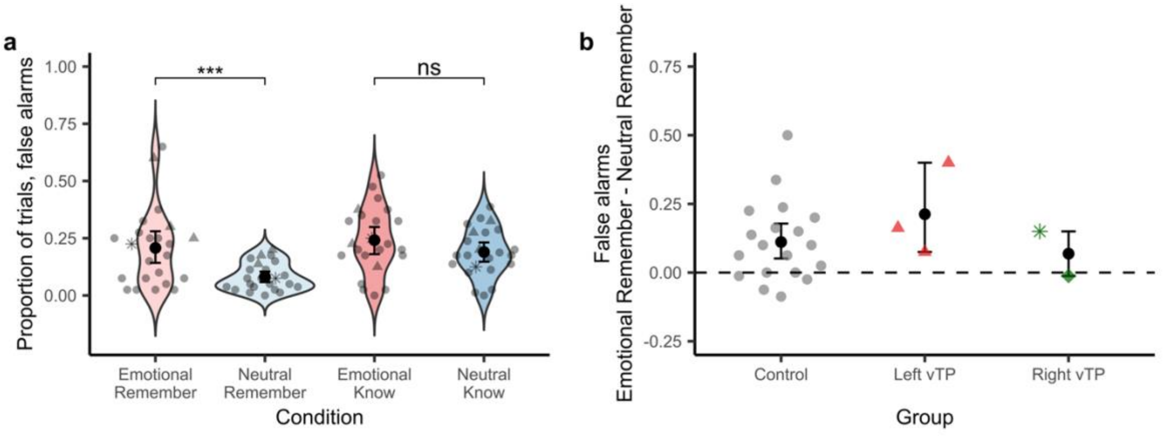


**Extended Data Fig. 7 | False alarms in the visual task. a**, The proportion of responses to new images which are false alarms shown for emotionally aversive and neutral pictures as a function of response, (Remember/Know). Scatter dots indicate individual means in proportion false alarms (• control patient, ▲ unilateral left vTP lesion, * bilateral vTP lesion). The black dots represent the group means of proportion false alarms and the error bars displays the 95% confidence interval of the mean. Significance levels: *** < .001 and ns = not significant. **b,** Descriptive data of false alarms displayed for the control group, the left ventral temporal pole group and the right ventral temporal pole group (▲ left vTP lesion, ♦ right vTP lesion, * bilateral vTP lesions, • control patient). The y-axis plots increased false alarm rates for aversive pictures compared to neutral pictures, focusing on remembered responses. A score above 0 indicates that the proportion false alarms was higher for aversive pictures than for neutral pictures. The data analysis of false alarms from the visual task is described in **Supplementary Data 4**.
