## Supplementary Information for "Emotional amnesia in humans with focal temporal pole lesions"

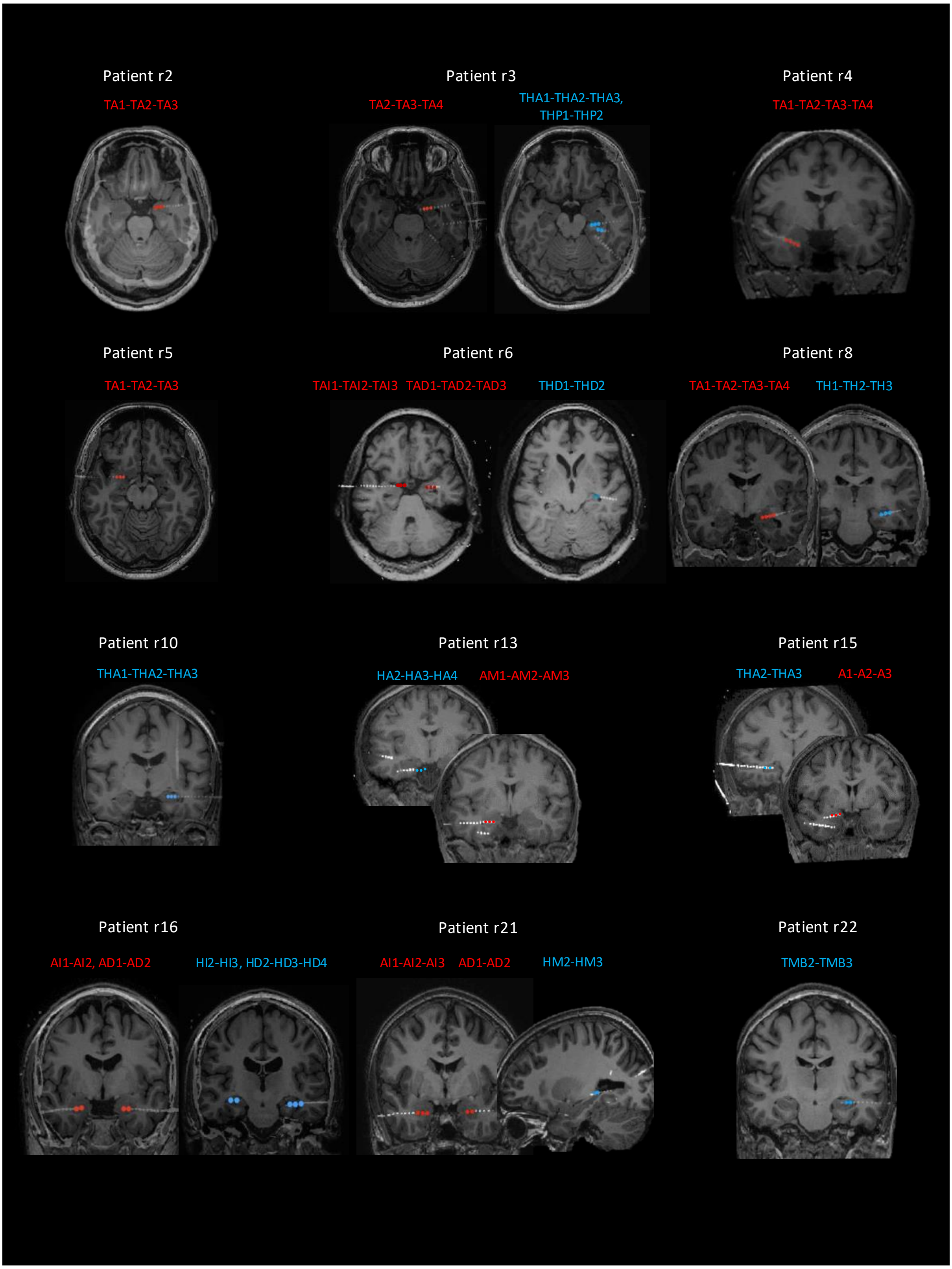
Patient r2
TA1-TA2-TA3
Patient r3
TA2-TA3-TA4
THA1-THA2-THA3,
THP1-THP2
Patient r4
TA1-TA2-TA3-TA4
Patient r5
TA1-TA2-TA3
Patient r6
TAI1-TAI2-TAI3
TAD1-TAD2-TAD3
THD1-THD2
Patient r8
TA1-TA2-TA3-TA4
TH1-TH2-TH3
Patient r10
THA1-THA2-THA3
Patient r13
HA2-HA3-HA4
AM1-AM2-AM3
Patient r15
THA2-THA3
A1-A2-A3
Patient r16
AI1-AI2, AD1-AD2
HI2-HI3, HD2-HD3-HD4
Patient r21
AI1-AI2-AI3
AD1-AD2
HM2-HM3
Patient r22
TMB2-TMB3

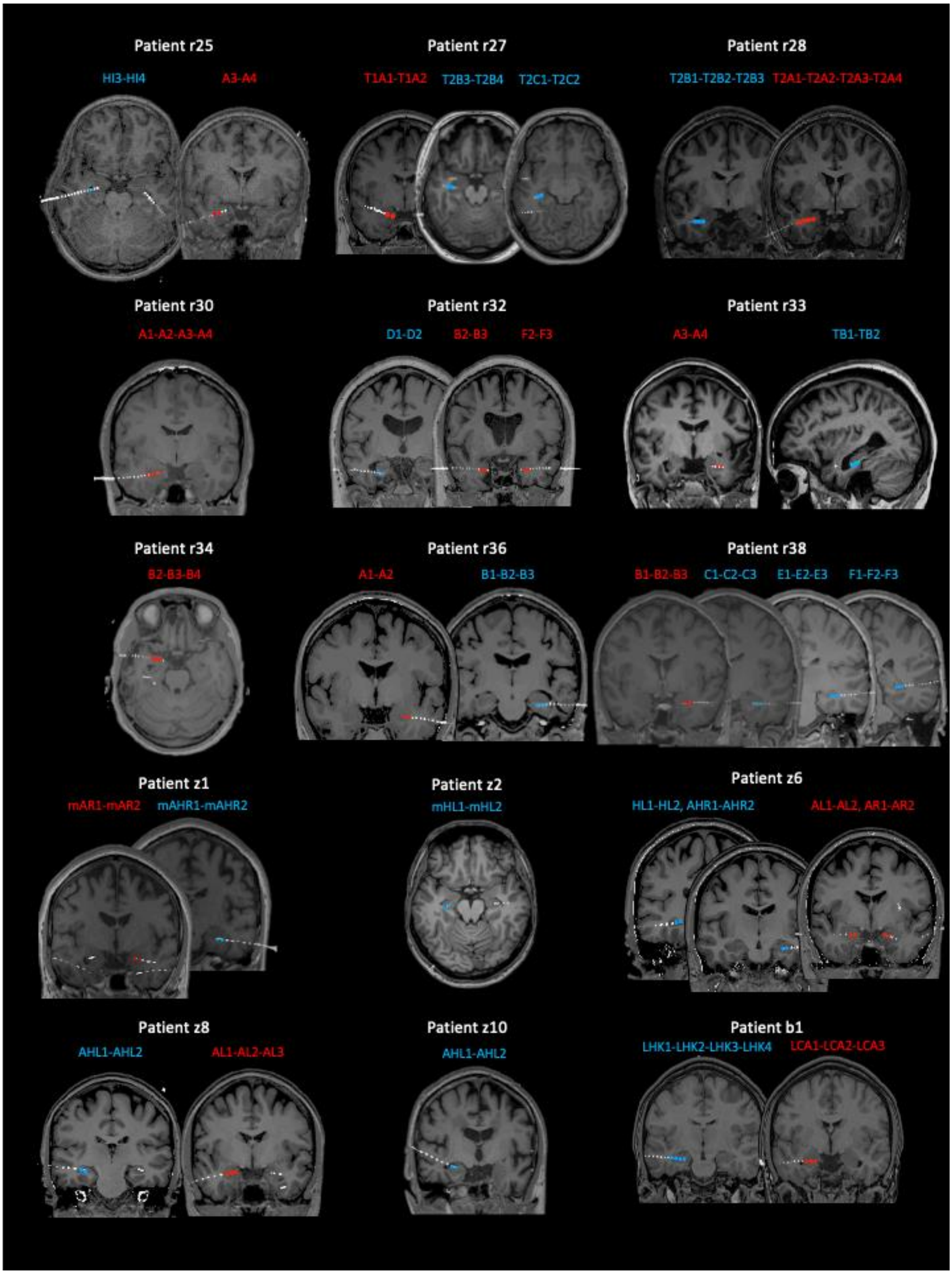


**Supplementary Figure 1. | Contacts used in the analyses displayed on post-operational CTs overlayed on pre-operational MRIs.** Red = amygdala contacts, blue = hippocampal contacts.

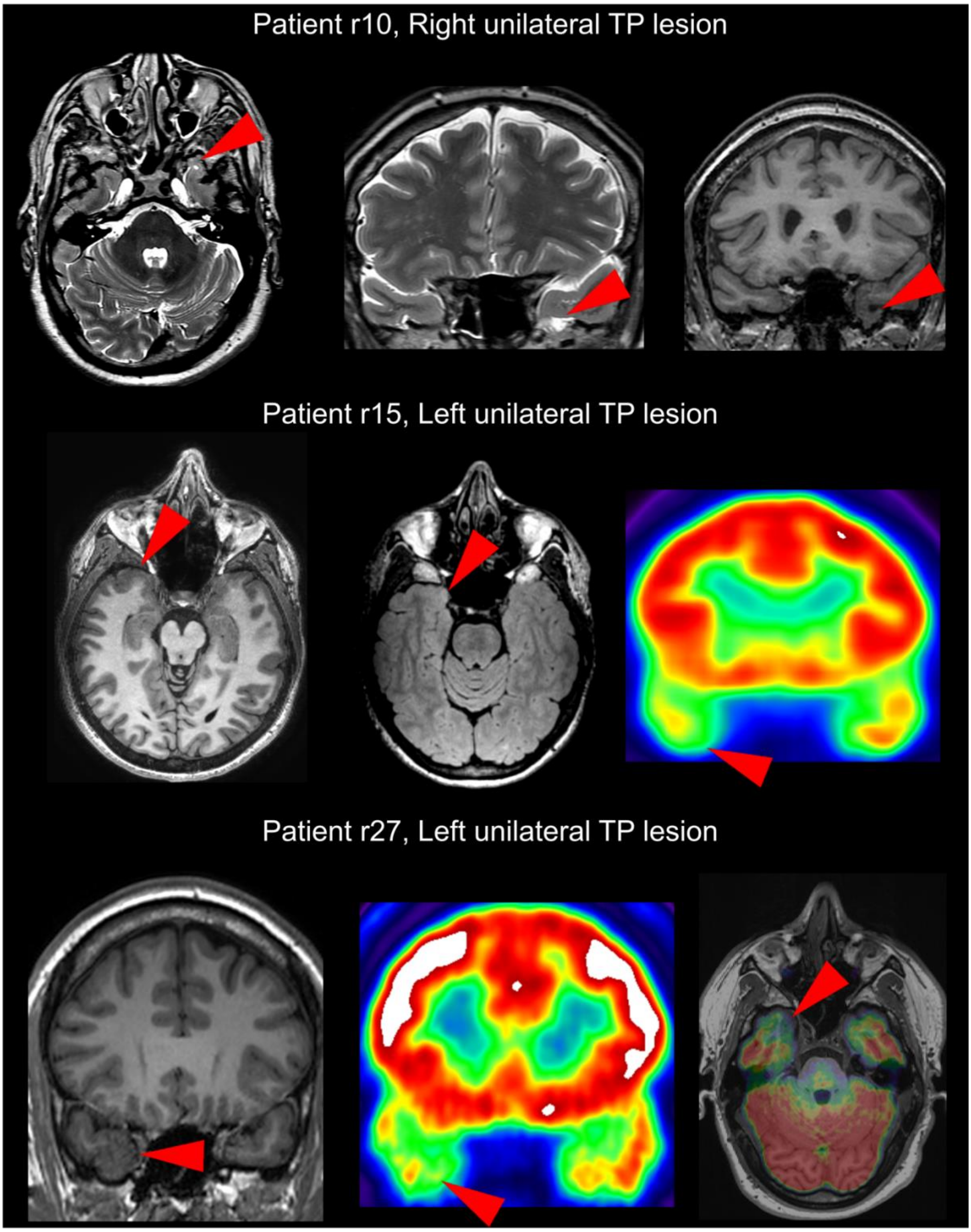
Patient r10, Right unilateral TP lesion
Patient r15, Left unilateral TP lesion
Patient r27, Left unilateral TP lesion

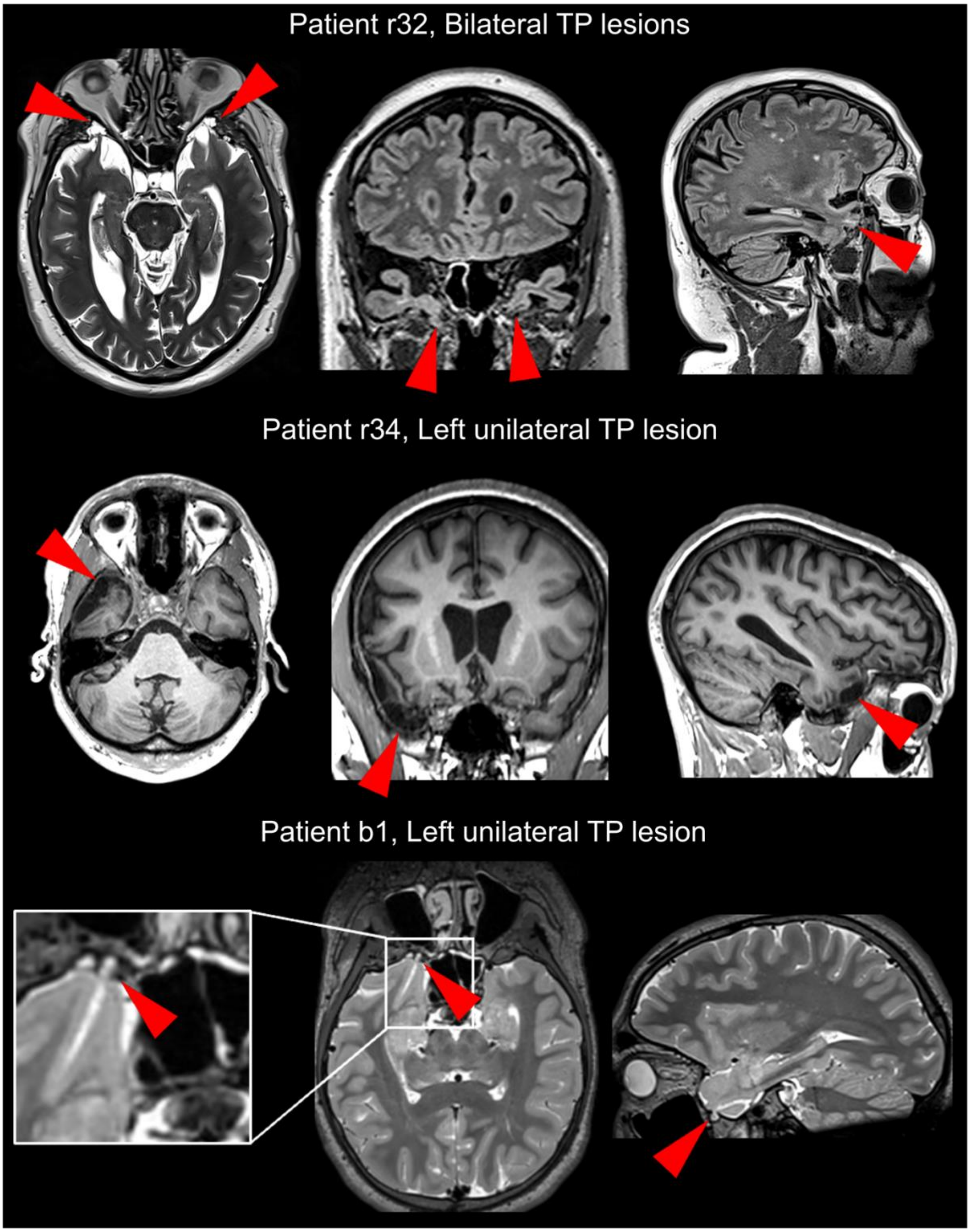


**Supplementary Figure 2. | MRI and PET images of patient's TP lesions.** Red arrow heads point to abnormality. r = patient from Ruber international hospital, Madrid, Spain. b = patient from Mara Hospital in Bielefeld, Germany.

**Supplementary Table 1. | Patient information.** Abbreviations: R = Ruber internacional hospital Madrid, b = Mara hospital Bielefeld, z = Swiss Epilepsy Centre Zürich. M = Male, F = Female, R = right, L = left, Am = amygdala, Hc = hippocampus, TP = temporal pole.

| ID | Age | Gender | Handedness | Aetiology | Lesion Location | Tasks Performed | iEEG regions | Group | Completed lists in verbal task |
|---|---|---|---|---|---|---|---|---|---|
| r1 | 26 | M | R | Polymicrogyria | Right perinsular region | Verbal | Only behaviour | Control | 30 |
| r2 | 38 | F | R | Hippocampal sclerosis plus focal dysplasia | Right hippocampal sclerosis plus porencephalic cyst over the parieto-occipital junction | Verbal + visual | Am | Control | 30 |
| r3 | 55 | M | R | Focal cortical dysplasia | Right temporal neocortex | Verbal | Am + Hc | Control | 30 |
| r4 | 21 | F | R | Focal cortical dysplasia plus gliosis | Extensive lesion over the left frontal region involving dorsolateral and orbitofrontal cortex and anterior border of the cingulum | Verbal + visual | Am | Control | 30 |
| r5 | 29 | M | R | Periventricular heterotopia plus focal cortical dysplasia | Bilateral occipital horn heterotopia plus focal dysplasia over the left occipital cortex | Verbal + visual | Am (verbal task) | Control | 30 |
| r6 | 49 | M | R | Hippocampal sclerosis | Left hippocampal sclerosis | Visual | Am + Hc | Control | NA |
| r8 | 28 | F | R | Focal cortical dysplasia | Extensive right posterior dysplasia involving the convexity and medial aspect of parietal, occipital and posterior temporal lobes. | Verbal | Am + Hc | Control | 30 |
| r10 | 59 | M | R | Encephalocele | Right anterior temporal pole and basal area. | Verbal | Only behaviour | Right TP | 30 |
| r12 | 28 | F | R | Focal cortical dysplasia | Right parietal-temporal region | Verbal | Only behaviour | Control | 30 |
| r13 | 42 | F | R | Focal cortical dysplasia | Right basal temporal cortex | Visual | Am + Hc | Control | NA |
| r15 | 35 | F | R | Focal cortical dysplasia | Left temporal pole | Verbal + visual | Am + Hc | Left TP | 22 |
| r16 | 30 | M | R | Reactive gliosis, diffuse | Medial wall of the left | Verbal + visual | Am + Hc | Control | 30 |

| | | | | | | | | | |
|---|---|---|---|---|---|---|---|---|---|
| | | | | microglia activation and small vessel vasculopathy | parietal region (precuneus and posterior cingulum) | | | | |
| r21 | 31 | F | R | Periventricular heterotopia | Left occipital horn heterotopia | Verbal + visual | Am + Hc (verbal task) | Control | 30 |
| r22 | 45 | F | R | Focal cortical dysplasia | main epileptic zone in right frontal operculum | Verbal + visual | Hc (verbal task) | Control | 30 |
| r25 | 29 | M | R | Focal cortical dysplasia | Right posterior temporobasal region | Verbal + visual | Am + Hc | Control | 30 |
| r27 | 25 | M | R | Focal cortical dysplasia | Left temporal pole | Verbal + visual | Am + Hc (verbal task) | Left TP | 30 |
| r28 | 25 | M | R | Focal cortical dysplasia | Left temporal lateral region | Verbal | Am + Hc (verbal task) | Control | 30 |
| r30 | 33 | F | R | Not operated (possible focal cortical dysplasia) | Left occipital medial region | Verbal | Am + Hc | Control | 15 |
| r32 | 59 | M | R | Encephalocele plus gliosis | Bilateral temporal pole | Verbal + visual | Am + Hc | Left TP (bilateral TP lesion) | 19 |
| r33 | 22 | F | R | Inflammatory lesion | Right parietal cortex | Verbal + visual | Am + Hc | Control | 22 |
| r34 | 18 | M | R | Postraumatic | Left temporopolar and temporolateral | Verbal + visual | Am | Left TP | 25 |
| r36 | 28 | F | R | Gliosis (posthemorraghe) | Right temporo-occipital region | Verbal + visual | Am + Hc | Control | 30 |
| r37 | 24 | F | L | Not operated (possible focal cortical dysplasia) | Left posterior temporobasal region | Visual | Only behaviour | Control | 30 |
| r38 | 18 | M | R | Pathology (Blumcke): Temporal neocortex with glial scar | Right temporal lateral region | Verbal + visual | Am + Hc | Control | 30 |
| b1 | 15 | M | R | (Meningo-) Encephalocele | Left basal temperal pole | Verbal | Am + Hc | Left TP | 34 |
| z1 | 51 | F | R | Hippocampal sclerosis | Left hippocampus | Visual | Hc | Control | NA |
| z2 | 37 | M | R | Focal cortical dysplacia | Right frontal cortex | Visual | Hc | Control | NA |
| z5 | 30 | M | R | Dysembroplastic neuroepithelia tumor | Left anterior hippocampus | Visual | Only behaviour | Control | NA |
| z6 | 29 | F | R | Unclear ethiology | Unclear | Visual | Am + Hc | Control | NA |
| z8 | 30 | F | R | Non-lesional | | Visual | Am + Hc | Control | NA |
| z10 | 56 | M | R | Hippocampal sclerosis | Right hippocampus | Visual | Hc | Control | NA |
| z12 | 24 | F | R | Hippocampal sclerosis | Right hippocampus | Visual | Only behaviour | Control | NA |

**Supplementary Table 2. | Sample in the behavioural data analyses in the verbal task.**

| | Patient ID | Group |
|---|---|---|
| 1 | r1 | Control |
| 2 | r2 | Control |
| 3 | r3 | Control |
| 4 | r4 | Control |
| 5 | r5 | Control |
| 6 | r8 | Control |
| 7 | r12 | Control |
| 8 | r15 | Left vTP |
| 9 | r16 | Control |
| 10 | r21 | Control |
| 11 | r22 | Control |
| 12 | r25 | Control |
| 13 | r27 | Left vTP |
| 14 | r28 | Control |
| 15 | r30 | Control |
| 16 | r32 | Left vTP (bilateral lesions) |
| 17 | r33 | Control |
| 18 | r34 | Left vTP |
| 19 | r36 | Control |
| 20 | r38 | Control |
| 21 | b1 | Left vTP |

**Supplementary Table 3. | Sample in the amygdala iEEG analyses in the verbal task**

| | Patient ID | Group | iEEG hemisphere |
|---|---|---|---|
| 1 | r2 | Control | Right |
| 2 | r3 | Control | Right |
| 3 | r4 | Control | Left |
| 4 | r5 | Control | Left |
| 5 | r8 | Control | Right |
| 6 | r15 | Left vTP | Left |
| 7 | r16 | Control | Bilateral |
| 8 | r21 | Control | Bilateral |
| 9 | r25 | Control | Left |
| 10 | r27 | Left vTP | Left |
| 11 | r28 | Control | Left |
| 12 | r30 | Control | Left |
| 13 | r32 | Left vTP (bilateral lesions) | Bilateral |
| 14 | r33 | Control | Right |
| 15 | r34 | Left vTP | Left |
| 16 | r36 | Control | Right |
| 17 | b1 | Left vTP | Left |

**Supplementary Table 4. | Sample in the hippocampus iEEG analyses in the verbal task.**

| | Patient ID | Group | iEEG hemisphere |
|---|---|---|---|
| 1 | r3 | Control | Right |
| 2 | r8 | Control | Right |
| 3 | r15 | Left vTP | Left |
| 4 | r16 | Control | Bilateral |
| 5 | r21 | Control | Left |
| 6 | r22 | Control | Right |
| 7 | r25 | Control | Left |
| 8 | r27 | Left vTP | Left |
| 9 | r28 | Control | Left |
| 10 | r32 | Left vTP (bilateral lesions) | Left |
| 11 | r33 | Control | Right |
| 12 | r36 | Control | Right |
| 13 | r38 | Control | Right |
| 14 | b1 | Left vTP | Left |

**Supplementary Table 5. | Sample in behavioural data analyses in the visual task**

| | Patient ID | Group |
|---|---|---|
| 1 | r2 | Control |
| 2 | r4 | Control |
| 3 | r5 | Control |
| 4 | r6 | Control |
| 5 | r13 | Control |
| 6 | r15 | Left vTP |
| 7 | r16 | Control |
| 8 | r21 | Control |
| 9 | r25 | Control |
| 10 | r27 | Left vTP |
| 11 | r32 | Left vTP |
| 12 | r33 | Control |
| 13 | r34 | Left vTP |
| 14 | r36 | Control |
| 15 | r37 | Control |
| 16 | r38 | Control |
| 17 | z1 | Control |
| 18 | z2 | Control |
| 19 | z5 | Control |
| 20 | z6 | Control |
| 21 | z8 | Control |
| 22 | z10 | Control |
| 23 | z12 | Control |

**Supplementary Table 6. | Sample in amygdala iEEG data analyses in the visual task.**

| | Patient ID | Group | iEEG hemisphere |
|---|---|---|---|
| 1 | r2 | Control | Right |
| 2 | r4 | Control | Left |
| 3 | r6 | Control | Bilateral |
| 4 | r13 | Control | Left |
| 5 | r15 | Left vTP | Left |
| 6 | r16 | Control | Bilateral |
| 7 | r21 | Control | Bilateral |
| 8 | r25 | Control | Left |
| 9 | r27 | Left vTP | Left |
| 10 | r32 | Left vTP (bilateral lesions) | Bilateral |
| 11 | r33 | Control | Right |
| 12 | r34 | Left vTP | Left |
| 13 | r36 | Control | Right |
| 14 | r38 | Control | Right |
| 15 | z6 | Control | Bilateral |
| 16 | z8 | Control | Left |

**Supplementary Table 7. | Sample in hippocampus iEEG data analyses in the visual task**

| | Patient ID | Group | iEEG hemisphere |
|---|---|---|---|
| 1 | r6 | Control | Right |
| 2 | r13 | Control | Left |
| 3 | r15 | Left vTP | Left |
| 4 | r16 | Control | Bilateral |
| 5 | r25 | Control | Left |
| 6 | r32 | Left vTP (bilateral lesions) | Left |
| 7 | r33 | Control | Right |
| 8 | r36 | Control | Right |
| 9 | r38 | Control | Right |
| 10 | z1 | Control | Right |
| 11 | z2 | Control | Right |
| 12 | z6 | Control | Bilateral |
| 13 | z8 | Control | Left |
| 14 | z10 | Control | Left |

**Supplementary Data 1 | Reaction time analysis of encoding data from the verbal task**

Reaction times (RTs) were entered into a GLMM with fixed effects of Group (control vs left ventral temporal pole) and Condition (emotional oddball vs perceptual oddball vs control words). There was a significant main effect of Condition ($\chi^2(2) = 14.58$, $P < .001$), while there was no main effect of Group ($\chi^2(1) = .70$, $P = .404$) and, importantly, no Group × Condition interaction ($\chi^2(2) = 1.04$, $P = .594$). Follow-up pairwise comparisons showed that RTs were slower for emotional oddballs (left vTP lesioned group: *m* = 1008ms, *SD* = 338ms; Control group: *m* = 1048, *SD* = 360) compared with control words (left vTP lesioned group: *m* = 951ms, *SD* = 288ms; Control group: *m* = 974ms, *SD* = 315ms; *z* = 3.64, *P* < .001) with responses being approximately 7% longer (*M* ratio = 1.07). RTs to perceptual oddballs were also slower than control words (left vTP lesioned group: *m* = 989ms, *SD* = 292ms; Control group: *m* = 1003ms, *SD* = 336ms, *z* = 2.31, *P* = .032) with around 4% longer RTs (*M* ratio = 1.04), while there was no difference in reaction times between emotional and perceptual oddballs (*z* = 1.11, *P* = .266, *M* ratio = 1.02).

**Supplementary Data 2 | vTP lesions do not affect reaction times to negative visual stimuli**

For the visual task, RTs to indoor/outdoor judgements during the encoding phase were analysed with a Group (control vs left vTP) × Emotion (negative vs neutral) mixed ANOVA. There was a main effect of Emotion (**Extended Data Fig. 4a**), reflecting slower RTs to negative compared to neutral images ($F(1,21) = 5.20$, $p = .033$, $\eta_p^2 = .198$). There was no main effect of Group ($F(1,21) = .85$, $p = .366$, $\eta_p^2 = .039$) and no Group × Emotion interaction ($F(1,21) = 1.43$, $p = .244$, $\eta_p^2 = .064$), indicating that left vTP lesions did not affect RTs to negative visual stimuli. Both the unilateral right vTP lesioned patient and the bilateral vTP lesioned patient showed slower RTs to emotional vs. neutral images, indicating that they experienced the emotionality of the images (mean difference between negative and neutral, right unilateral TP patient = 579ms, $z$ compared with the control group = 3.49; bilateral vTP patient = 125ms, $z$ = .04).

Next, RTs during the recognition test of the visual task were analysed using a Group (control vs left vTP) × Emotion (negative vs neutral) × Memory (remember vs know vs correct rejection) mixed ANOVA. One control patient (z2) was excluded for not responding *know* to any emotional stimuli. A main effect of Memory was observed ($F(1,20) = 10.18$, $p < .001$, $\eta_p^2 = .337$). Follow-up pairwise comparisons showed faster RTs for remember responses compared to both *correct rejections* ($t(20) = -2.90$, $p = .034$) and *know* responses ($t(20) = -3.90$, $p = .003$) and faster RTs for *correct rejections* than *know* responses ($t(20) = -2.63$, $p = .024$). Importantly, there was no interaction involving the factor Group. Thus, RTs to visual stimuli were normal in the left vTP patients. The bilateral vTP lesioned patient followed the same pattern as the control group with fastest RTs for *remember* responses followed by *correct rejections* and *know* responses (*mean* RTs; *remember* = 1452 ms, *correct rejections* = 1671 ms, *know* = 1792 ms) while the right vTP lesioned patient showed fastest RTs for *correct rejections* ($m$ = 2037ms), followed by *remember* responses ($m$ = 2452 ms) and *know* responses ($m$ = 2725 ms). Given that the right vTP patients had fewer correct *remember* responses to emotional items compared to neutral items we describe these patients' RTs for these conditions even though there was no Emotion x Memory interaction in the omnibus ANOVA for RTs described above. Mirroring the accuracy results, there was a slowing in RTs for *remember* responses to emotional images compared with *remember* responses to neutral images in both the unilateral right vTP lesioned patient (mean difference = 356ms, $z$ compared with control group = 1.07) and in the bilateral vTP lesioned patient (mean difference = 63ms, $z$ = .45).

**Supplementary Data 3 | Left temporal pole lesions do not disrupt hippocampus iEEG responses**

We first investigated hippocampal responses to salience in the verbal task. Similar to the amygdala, there was no significant difference in power between emotionally negative words and perceptual oddballs, so the data was collapsed over both stimulus types. Previous research has shown increases in theta and gamma power in the hippocampus for unexpected oddball stimuli compared to expected stimuli[1]. We observed an increase in theta power (6-8Hz) for oddballs compared to control words between approximately 910-1500ms after stimulus presentation ($n$ = 14, $t_{sum}$ = 494.51, $P$ = .016 cluster corrected; **Extended Data Fig. 5a and 5b**). There was no evidence that this theta effect was lateralized (left $n$ = 8, right $n$ = 7, $t(13)$ = -1.73, $p$ = .108); **Extended Data Fig. 5c**) and there was no difference between the left temporal pole group and the control group (control group $n$ = 4, left vTP group $n$ = 4, $t(6)$ = 1.03, $p$ = .341, **Extended Data Fig. 5d**).

Next, we investigated hippocampal responses to emotionally negative images in the visual task. There was a main effect of Emotion in the hippocampus ($n$ = 14, $t_{sum}$ = 1397.50, $P$ = .017 cluster corrected), with emotional pictures being related to higher gamma power compared to neutral pictures between 510-1320ms and between 37-65Hz (**Extended Data Fig. 5e and 5f**). There was no evidence of lateralization for this effect (left $n$ = 8, right $n$ = 8, $t(14)$ = -.29, $p$ = .776, **Extended Data Fig. 5g**). The two patients with left TP lesions both had higher gamma power for emotional than neutral pictures in this cluster, suggesting that they had a normal response (in the control group, 10 out of 12 patients showed the same pattern, **Extended Data Fig. 5h**).

We then examined hippocampus responses related to subsequent memory during encoding in both the verbal and the visual task. Successful encoding has been related both decreases and increases in theta power[2–7], to decreases in alpha and beta power[3,5,7] and to increases in gamma power[3–6,8–10]. For the verbal task, we excluded the four first words from each list to control for primacy effects. Subsequently remembered words were related to a reduction in beta power (15-22Hz) around 810-1160ms after the presentation of the word stimulus (*n* = 14, $t_{sum}$ = -473.81, *P* = .027 cluster corrected; **Extended Data Fig.6a and 6b**). The effect was not lateralized (*t*(13) = -.22, *p* = .833; **Extended Data Fig.6c**) and not different between the left vTP group (*n*= 4) and the control group (*n* = 4; *t*(6) = -1.0, *p* = .358; **Extended Data Fig.6d**). In the visual task, there was a reduction in alpha/beta power (8-14Hz) for subsequently remembered compared to forgotten images from around 680-1340ms after stimulus presentation (*n* = 14, $t_{sum}$ = -497.78, *P* = .028 cluster corrected; **Extended Data Fig.6e and 6f**). The effect was larger in the right (*n* = 8) than in the left (*n* = 8) hemisphere (*t*(14) = 3.00, *p* = .010, *d* = 1.50; **Extended Data Fig.6g**). The alpha/beta effect was not significant in the left hippocampus (*t*(7) = -.44, *p* = .674), so it was not possible to investigate if the two left vTP patients had a normal response as they only have contacts in the left hippocampus (**Extended Data Fig.6h**). There was also a main effect of subsequent memory (*n* = 14, $t_{sum}$ = 1149.14, *p* = .013) in the gamma band, with higher gamma power for remembered compared to forgotten pictures between 0-870ms and 40-67Hz (**Extended Data Fig.6i and 6j**). The early timing of this effect suggests that it may partly reflect preparatory processes followed by encoding related activity. This effect was not lateralized (*n* left = 8, *n* right = 8; *t*(14) = -.28, *p* = .785; **Extended Data Fig.6k**). One of the left vTP subjects had a normal response and one did not show a subsequent memory gamma effect. This effect was however also absent in one out of 6 control subjects (17%, **Extended Data Fig.6l**). Across all five hippocampus effects to saliency and subsequent memory, the data indicates that the left vTP group had normal hippocampus responses (not different from the control group). There were no subsequent memory effects in the theta band in the two tasks. The majority of the hippocampal contacts in this study were placed in the anterior hippocampus and previous research suggest that hippocampal theta subsequent memory effects are larger in the posterior hippocampus[11], so this may explain why no theta subsequent memory effects were observed.

**Supplementary Data 4 | Analysis of false alarms in the visual task**

False alarms in the recognition memory test in the visual were analysed using a Group (control vs left vTP) x Emotion (aversive vs neutral) x Response (remember vs know) mixed ANOVA. There was a main effect of Emotion showing that there were more false alarms to emotional than to neutral pictures ($F(1,21) = 14.45$, $P = .001$, $\eta_p^2 = .408$, **Extended Data Fig. 7a**). In addition, there was an Emotion x Memory interaction ($F(1,21) = 4.47$, $P = .047$, $\eta_p^2 = .176$). Follow-up pairwise comparisons showed that there were more false alarms for emotional than neutral images for *remember* responses ($t(21) = 3.85$, $p < .001$, $d = .80$) while there was no difference in false alarms between emotional and neutral images for *know* responses ($t(21) = 1.04$, $p = .309$, $d = .22$). There was a trend towards a main effect of group with left vTP patients having more false alarms than the control group overall ($F(1,21) = 3.19$, $P = .088$), but there were no interactions involving the factor Group (all $P$s $\geq .239$).

The two right vTP patients showed a numerical reduction in hits for emotional remembered responses compared to neutral remembered responses. We next investigated if this reduction in emotional memory was driven by a reduction in remember responses overall as would be indicated in reduced false alarms for emotional items compared to neutral items in this group. There was no evidence for this as the patient with bilateral lesions had more false alarms for emotionally negative than for neutral items (.15, $z$ compared to control group = .27) and the patient with a unilateral right vTP lesion had close to no difference in false alarms to aversive than to neutral items (-.01, $z$ compared with control group = -.85, **Extended Data Fig 7b**).
